# Unraveling the functions of uncharacterized transcription factors in *Escherichia coli* using ChIP-exo

**DOI:** 10.1101/2021.06.10.447994

**Authors:** Ye Gao, Hyun Gyu Lim, Hans Verkler, Richard Szubin, Daniel Quach, Irina Rodionova, Ke Chen, James T. Yurkovich, Byung-Kwan Cho, Bernhard O. Palsson

## Abstract

Bacteria regulate gene expression to adapt to changing environments through transcriptional regulatory networks (TRNs). Although extensively studied, no TRN is fully characterized since the identity and activity of all the transcriptional regulators that comprise a TRN are not known. Here, we experimentally evaluate 40 uncharacterized proteins in *Escherichia coli* K-12 MG1655, which were computationally predicted to be transcription factors (TFs). First, we used a multiplexed ChIP-exo assay to characterize genome-wide binding sites for these candidate TFs; 34 of them were found to be DNA-binding protein. We then compared the relative location between binding sites and RNA polymerase (RNAP). We found 48% (283/588) overlap between the TFs and RNAP. Finally, we used these data to infer potential functions for 10 of the 34 TFs with validated DNA binding sites and consensus binding motifs. These TFs were found to have various roles in regulating primary cellular processes in *E. coli*. Taken together, this study: (1) significantly expands the number of confirmed TFs, close to the estimated total of about 280 TFs; (2) predicts the putative functions of the newly discovered TFs, and (3) confirms the functions of representative TFs through mutant phenotypes.

## Introduction

Bacteria employ a broad range of mechanisms to regulate gene expression to achieve and maintain phenotypic states (1). One of these mechanisms relies on promoter recognition by the RNA polymerase (RNAP) and its subsequent initiation of transcription (2). However, the core enzyme is unable to recognize promoters or to initiate transcription without the assistance of one of a set of sigma factors. A sigma factor binds to the core enzyme, forming a complex known as RNA polymerase holoenzyme, which is able to orchestrate transcription initiation from specific promoters (1). Additionally, transcription factors (TFs) bind to intergenic regulatory regions, preventing or promoting RNAP binding upstream from a transcription start site (3). Thus, the identification of transcription factors and the transcription factor binding sites (TFBSs) is fundamental to understanding how an organism responds to varying phenotypic demands through transcriptional regulation.

Chromatin immunoprecipitation (ChIP) technologies have enabled the genome-wide characterization of TFBSs (4, 5). Comprehensively mapping these regulatory interactions is important for understanding how an organism achieves and maintains different phenotypic states (6). However, the current knowledge of the transcriptional regulatory network (TRN) for the well-studied model *Escherichia coli* K-12 MG1655 strain is still incomplete (7, 8). To address this challenge, we previously developed a pipeline for computational prediction followed by experimental validation of candidate TFs in *E. coli* (9). The initial use of this pipeline resulted in the characterization of ten novel TFs in *E. coli*, chosen from a rank-ordered list of computationally predicted candidates.

To achieve a comprehensive characterization of the *E. coli* K-12 MG1655 TRN, we employ this pipeline again in a high-throughput manner to characterize an additional 40 candidate TFs. We assembled a collection of data sets, including information about TFBSs and consensus DNA sequences of binding motifs, relative location of RNAP holoenzyme and TFBSs in the promoter region, and gene expression profiling. We integrated these diverse data to classify candidate TFs into three different groups—type I global regulator (>100 target genes, N=1), type II local regulator (2-100 target genes, N=29), and type III single-target regulator (N=4) (10) — and to explore the putative functions of 10 validated TFs with detailed analysis. These results illustrate that candidate TFs have a varied number of regulatory targets and participate in the primary cellular processes in *E. coli*, from replication, transcription, and nutrition metabolism, to stress responses. Taken together, our results expand the total number of *E. coli* TFs to 276 (an increase of ∼12%), and support the estimated total of 280∼300 TFs comprising the TRN in *E. coli* K-12 MG1655 (11).

## Methods and Materials

### Computational prediction of candidate TFs

We previously generated a list of candidate TFs and 16 of the top candidates were used to assess the discovery pipeline (9). Ten of the 16 candidates were found to be TFs. Here, we extended the experimental validation of these computationally predicted targets by selecting additional candidates from this previously generated list. Briefly, the list was generated using the TFpredict algorithm (12) modified for use with bacterial genomes (9). The TFpredict algorithm takes a protein sequence as input and generates a quantified score in the range [0,1] that represents the likelihood of that protein being a TF based on sequence homology, where a score of 1 represents the highest confidence. We selected 40 of the top candidate TFs from this rank-ordered list. See (9) for a full description of the computational methods.

### Bacterial strains, media, and growth conditions

All strains used in this study are *E. coli* K-12 MG1655 and its derivatives, deletion strains, and myc-tagged strains (Dataset S1). For ChIP-exo experiments, the *E. coli* strains harboring 8-myc were generated by a λ red-mediated site-specific recombination system targeting the C-terminal region as described previously (13). For ChIP-exo experiments, glycerol stocks of *E. coli* strains were inoculated into M9 minimal media (47.8 mM Na_2_HPO_4_, 22 mM KH_2_PO_4_, 8.6 mM NaCl, 18.7 mM NH_4_Cl, 2 mM MgSO_4_, and 0.1 mM CaCl_2_) with 0.2% (w/v) glucose. M9 minimal media was also supplemented with 1 mL trace element solution (100X) containing 1 g EDTA, 29 mg ZnSO_4_.7H_2_O, 198 mg MnCl_2_.4H_2_O, 254 mg CoCl_2_.6H_2_O, 13.4 mg CuCl_2_, and 147 mg CaCl_2_ per liter. The culture was incubated at 37 °C overnight with agitation and then was used to inoculate the fresh media (1/200 dilution). The volume of the fresh media was 150 mL for each biological replicate. The fresh culture was incubated at 37 °C with agitation to the mid-log phase (OD_600_ ≈ 0.5). To create oxidative stress, the overnight cultures were inoculated to an optical density OD_600_ = 0.01 into the fresh 70 mL of Glucose M9 minimal medium in a 500-mL flask supplemented with 250 μM paraquat (PQ) at OD_600_ = 0.3 and incubated for 20 min with stirring (**Dataset S2**).

To address the susceptibility of bacterial cells to H_2_O_2_, mid-log phase cells were treated with 60 mM H_2_O_2_ for 15 min (cells were washed by TBS before treatment). Serial dilutions were then prepared, and 10 μl of aliquots from the dilutions were spotted in triplicate on plates and incubated at 37°C overnight. The sensitivity of cells to the lethal effects of the stressor was expressed as percent survival of treated cells relative to that of untreated cells determined at the time of treatment.

The effects of carbon sources on cell growth were examined by growing *E. coli* K-12 MG1655 and *yciT* deletion strains on different carbon sources: (1) M9 minimal glucose (0.2% w/v) medium; (2) M9 minimal fructose (0.2% w/v) medium; and (3) M9 minimal sorbitol (0.2% w/v) medium. Cells grown overnight on M9 minimal medium with one sole carbon source (glucose, fructose, or sorbitol) at 37°C with agitation were inoculated into these three kinds of fresh media, then were intubated at 37°C with agitation. Similarly, the effects of the osmotic stress on cell growth were examined by growing *E. coli* K-12 MG1655 and *yciT* deletion strains on M9 minimal sorbitol (0.2% w/v) medium with or without 0.5 M NaCl. Cells grown overnight on M9 minimal medium with 0.2% sorbitol at 37°C with agitation were inoculated into fresh media with sorbitol (0.2% w/v) with or without 0.5 M NaCl. All of the growth curves were measured every 30 min by six independent experiments at least and recorded by OD_600_ using a Bioscreen C (Growth curves USA).

### Multiplexed ChIP-exo experiment

A multiplexed ChIP-exo experiment was performed on a simple modification of our standard ChIP-exo method described previously (14). Here, after ligating the first adapter to each sample separately, the samples are then pooled together and this pool undergoes the remainder of the enzymatic reactions used for library preparation. Each sample receives a different first adapter bearing a unique 6-base sequence (barcode) which allows for the data to be demultiplexed. To identify each candidate TF binding maps *in vivo*, the DNA bound to each candidate TF from formaldehyde cross-linked *E. coli* cells were isolated by chromatin immunoprecipitation (ChIP) with the specific antibodies that specifically recognize myc tag (9E10, Santa Cruz Biotechnology), and Dynabeads Pan Mouse IgG magnetic beads (Invitrogen) followed by stringent washings as described previously (15). Cells were grown in glucose minimal medium to OD_600_ = 0.5, and incubated with 1% formaldehyde (Thermo Scientific) for 25 min at room temperature. Formaldehyde was quenched by 2.5 M glycine (Thermo Fisher Scientific) for an additional 5 min and the cells were washed with ice-cold TBS (Thermo Fisher Scientific) three times, and lysed with Ready-lyse lysozyme solution (Epicentre). Lysates were sonicated using a sonicator (QSonic) to generate 300–500 bp randomly sheared chromosomal DNA fragments. The extent of shearing was monitored with a 1% agarose gel and confirmed by separation on a 2100 High sensitivity Bioanalyzer chip (Agilent Technologies) upon completion of the immunoprecipitation. Immunoprecipitation was carried out at 4°C with overnight incubation and 15 μl anti-c-myc mouse antibody (9E10, Santa Cruz Biotechnology). The protein of interest, together with its cross-linking DNA and covalently bound mouse antibody, was captured with 50 μl Dynabeads Pan mouse IgG (Invitrogen), washed by washing buffer I (50 mM Tris-HCl (pH 7.5), 140 mM NaCl, 1 mM EDTA,1% Triton X-100).

ChIP materials (chromatin-beads) were used to perform on-bead enzymatic reactions of the ChIP-exo method (5). Briefly, the sheared DNA of chromatin-beads was repaired by the NEBNext End Repair Module (New England Biolabs) followed by the addition of a single dA overhang and ligation of the first adaptor (5’-phosphorylated) using dA-Tailing Module (New England Biolabs) and NEBNext Quick Ligation Module (New England Biolabs), respectively. The first adaptor was designed to have different indices to distinguish different DNA samples after the sequencing. After ligation, multiple ChIP materials could be pooled together. Nick repair was performed by using PreCR Repair Mix (New England Biolabs). Lambda exonuclease- and RecJ_f_ exonuclease-treated chromatin was eluted from the beads and overnight incubation at 65 °C reversed the protein-DNA cross-link. RNAs- and Proteins-removed DNA samples were used to perform primer extension and second adaptor ligation with following modifications. The DNA samples incubated for primer extension as described previously (14) were treated with dA-Tailing Module (New England Biolabs) and NEBNext Quick Ligation Module (New England Biolabs) for second adaptor ligation. The DNA sample purified by GeneRead Size Selection Kit (Qiagen) was enriched by polymerase chain reaction (PCR) using Phusion High-Fidelity DNA Polymerase (New England Biolabs). The amplified DNA samples were purified again by GeneRead Size Selection Kit (Qiagen) and quantified using Qubit dsDNA HS Assay Kit (Life Technologies). Quality of the DNA sample was checked by running Agilent High Sensitivity DNA Kit using Agilent 2100 Bioanalyzer (Agilent) before sequenced using HiSeq 2500 (Illumina) following the manufacturer’s instructions. The antibody (NT63, Biolegend) that specifically recognizes RNA polymerase β was used to conduct the ChIP-exo experiment to detect the binding sites of RNA polymerase in *E. coli* K-12 MG1655. The antibody (2G10, Biolegend) that specifically recognizes σ^70^ was used to detect the binding sites of σ^70^ in *E. coli* K-12 MG1655. Each step was also performed following the manufacturer’s instructions. ChIP-exo experiments were performed in biological duplicates (Dataset S3 and S4).

### Peak calling for ChIP-exo dataset

Peak calling was performed as previously described (14). Sequence reads generated from ChIP-exo were mapped onto the reference genome (NC_000913.2) using bowtie (16) with default options to generate SAM output files. The MACE program was used to define peak candidates from biological duplicates for each experimental condition with sequence depth normalization (17). To reduce false-positive peaks, peaks with signal-to-noise (S/N) ratio less than 1.5 were removed. The noise level was set to the top 5% of signals at genomic positions because top 5% makes a background level in a plateau and top 5% intensities from each ChIP-exo replicates across conditions correlate well with the total number of reads (14, 18, 19). The calculation of S/N ratio resembles the way to calculate ChIP-chip peak intensity where IP signal was divided by Mock signal. Then, each peak was assigned to the target gene, according to genomic position (**Supplementary Figure 1**). Genome-scale data were visualized using MetaScope (https://sites.google.com/view/systemskimlab/software?authuser=0) and NimbleGen’s SignalMap software.

### Motif search from ChIP-exo peaks

The consensus DNA sequence motif analysis for validated TFs was performed using the MEME software suite (20). For YciT, YcjW, YdcN, YdhB, YfeC, YfeD, and YidZ, sequences in binding regions were extracted from the reference genome (NC_000913.2).

### COG functional enrichment

The regulons were categorized according to their annotated clusters of orthologous groups (COG) category (21). Functional enrichment of COG categories in the target genes was determined by performing a hypergeometric test, and *p*-value < 0.01 was considered significant.

### Transcriptomics

RNA-seq was performed using two biological replicates (Dataset S5). The strains were grown under the same conditions as those used in the ChIP-exo experiments. Transcripts were stabilized by mixing 3 mL of cell cultures at the mid-log phase with 6 mL of RNAprotect Bacteria Reagent (Qiagen). Samples were immediately vortexed for 5 sec, incubated for 5 min at room temperature, and then centrifuged at 5000 g for 10 min. The supernatant was decanted and any residual supernatant was removed by inverting the tube once onto a paper towel. Total RNA samples were then isolated using RNeasy Plus Mini kit (Qiagen) following the manufacturer’s instruction. Samples were then quantified using a NanoDrop 1000 spectrophotometer (Thermo Scientific) and quality of the isolated RNA was checked by running RNA 6000 Pico Kit using Agilent 2100 Bioanalyzer (Agilent). Paired-end, strand-specific RNA-seq library was prepared using KAPA RNA Hyper Prep kit (KAPA Biosystems), following the instructions (22, 23). Resulting libraries were analyzed on an Agilent Bioanalyzer DNA 1000 chip (Agilent). Sequencing was performed on a Hiseq 2500 sequencer at the Genomics Core facility of University of California, San Diego.

### Calculation of differentially expressed genes

Expression profiling was performed as previously described (14). Raw sequence reads generated from RNA-seq were mapped onto the reference genome (NC_000913.2) using bowtie v1.2.3 (16) with the maximum insert size of 1000 bp, and two maximum mismatches after trimming 3 bp at 3’ ends (16). Transcript abundance was quantified using summarizeOverlaps from the R GenomicAlignments package, with strand inversion for the dUTP protocol and strict intersection mode (24). We then calculated the dispersion and differential expression level of each gene using DESeq2 (25). In DESeq2, it uses empirical Bayes shrinkage for dispersion estimation, which substantially improves the stability and reproducibility of analysis results compared to maximum-likelihood-based solutions. This also makes DESeq2 applicable for small studies with few replicates (25). Transcripts per Million (TPM) were calculated by DESeq2. For significance testing, DESeq2 uses a Wald test to calculate *p*-value. The Wald test calculates *p*-values from the subset of genes that pass an independent filtering step, and they are adjusted for multiple testing using the procedure of Benjamini and Hochberg (25). Expression with log_2_ fold-change ≥ log_2_(2.0) and adjusted *p*-value < 0.05 or log_2_fold-change ≤ -log_2_(2.0) and adjusted *p*-value < 0.05 was considered as differentially expressed (Dataset S6).

### Structural analysis of candidate TFs

Homology models of the candidate transcription factors YidZ, YfeC, YciT, YcjW, YdcN and YgbI were constructed using the SWISS-MODEL pipeline (26). Multiple templates were analyzed, and inference of the oligomeric state was based on the reported interface conservation scores to existing complexes of similar sequence identity. The structures were annotated using information in UniProt (27) and visualized with VMD (28).

## Results

Here, we describe the discovery and characterization of candidate TFs in *E. coli* following our previously reported and validated pipeline (9). First, we present an overview of the genome-wide binding sites, determined by ChIP-exo, for these candidate TFs, highlighting their structural and functional properties. We then describe the local regulation of transcription initiation by these candidate TFs through a separate ChIP-exo screen for the RNAP holoenzyme. We then characterize the putative functions of 10 of the candidate TFs in *E. coli*, which provide insights into biological roles (**Figure 1**). Finally, three (YbcM, YciT, YgbI) of ten TFs were selected for mutant phenotype analysis.

**Figure 1.**
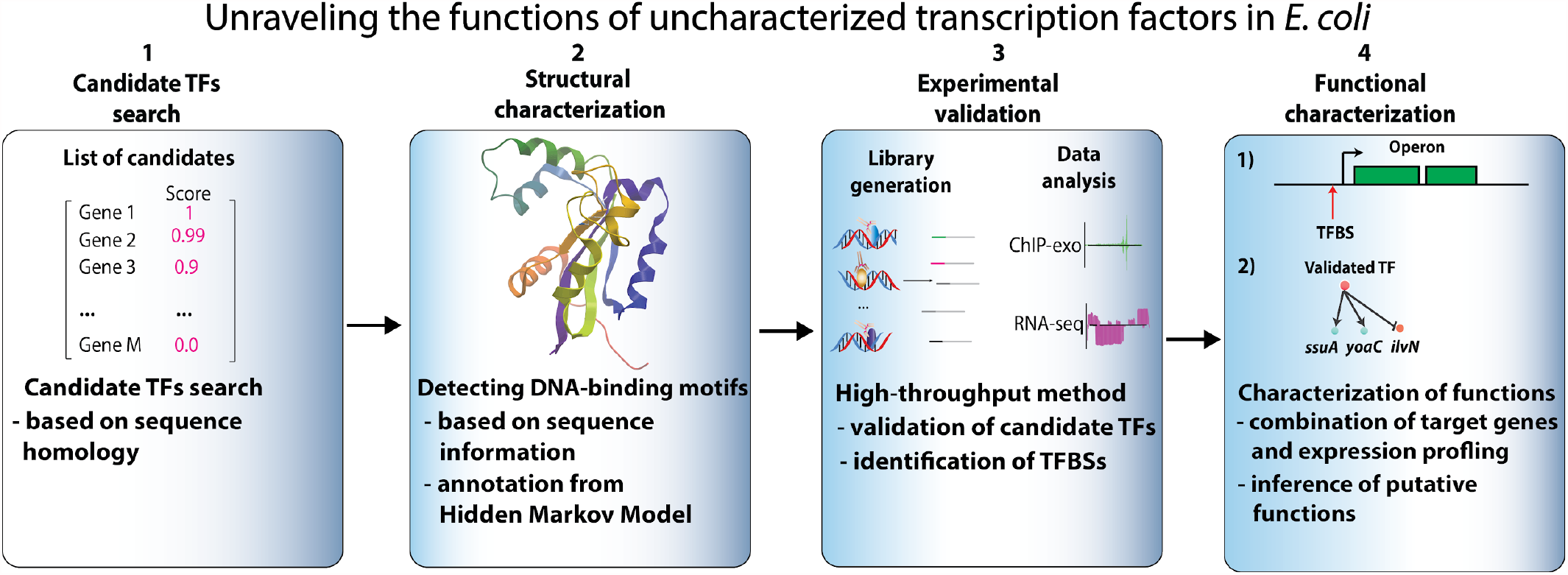
A systematic approach to identify and validate candidate transcription factors in *E. coli* K-12 MG1655. The approach used in this study can be divided into four steps: 1) We examined 40 computationally predicted candidate TFs from our previous study; 2) For each candidate TF, we highlighted its structural features based on the annotation from hidden Markov models; 3) We performed experimental validation using multiplexed ChIP-exo; and 4) We combined the genome-wide binding sites with expression profiling data to characterize regulatory roles of representative TFs with a suite of experimental tests.

### Putative transcription factors in *E. coli* K-12 MG1655

Previously, we had generated a rank-ordered list of candidate TFs from a list of uncharacterized genes (“y-genes”) using a homology-based algorithm (9). We experimentally tested 16 of the top hits from this list, verifying that ten (62.5%) were indeed TFs. In the present study, we selected an additional 40 candidate TFs from the list of candidates and experimentally tested them in a high-throughput manner (multiplexed ChIP-exo) (**Table 1**). In the time since we initiated this TF discovery effort, several of the candidate TFs have been independently suggested to be TFs using *in vitro* assays: ComR (YcfQ) (29), YcjW (30), SutR (YdcN) (31), RcdB (YhjC) (32), NimR (YeaM) (19), CsqR (YihW) (33, 34), YqhC (35). Our results provide *in vivo* genome-wide binding of these reported TFs, which is important for expanding the knowledge of the target genes and for building complete TRNs in *E. coli*.

**Table 1.**
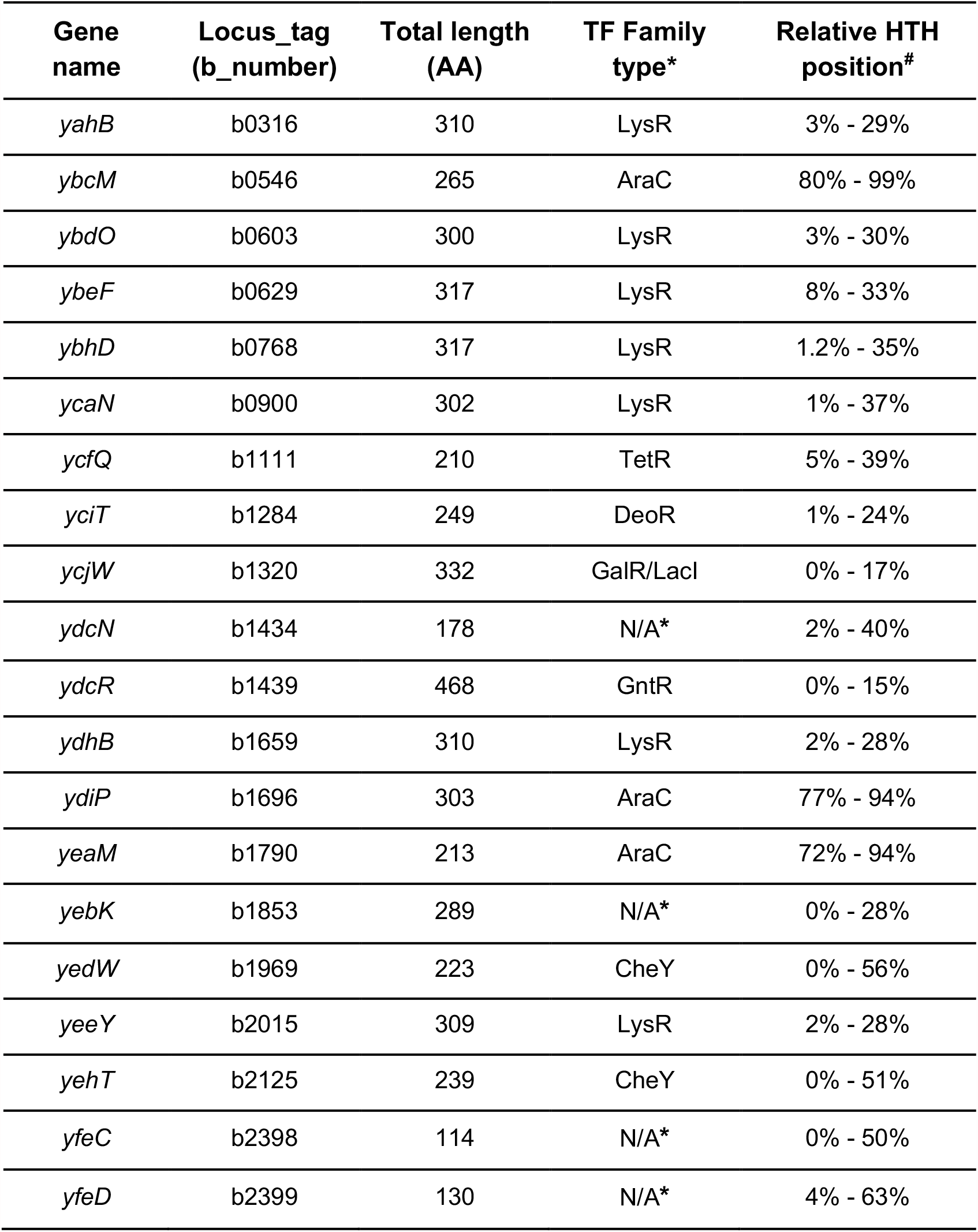
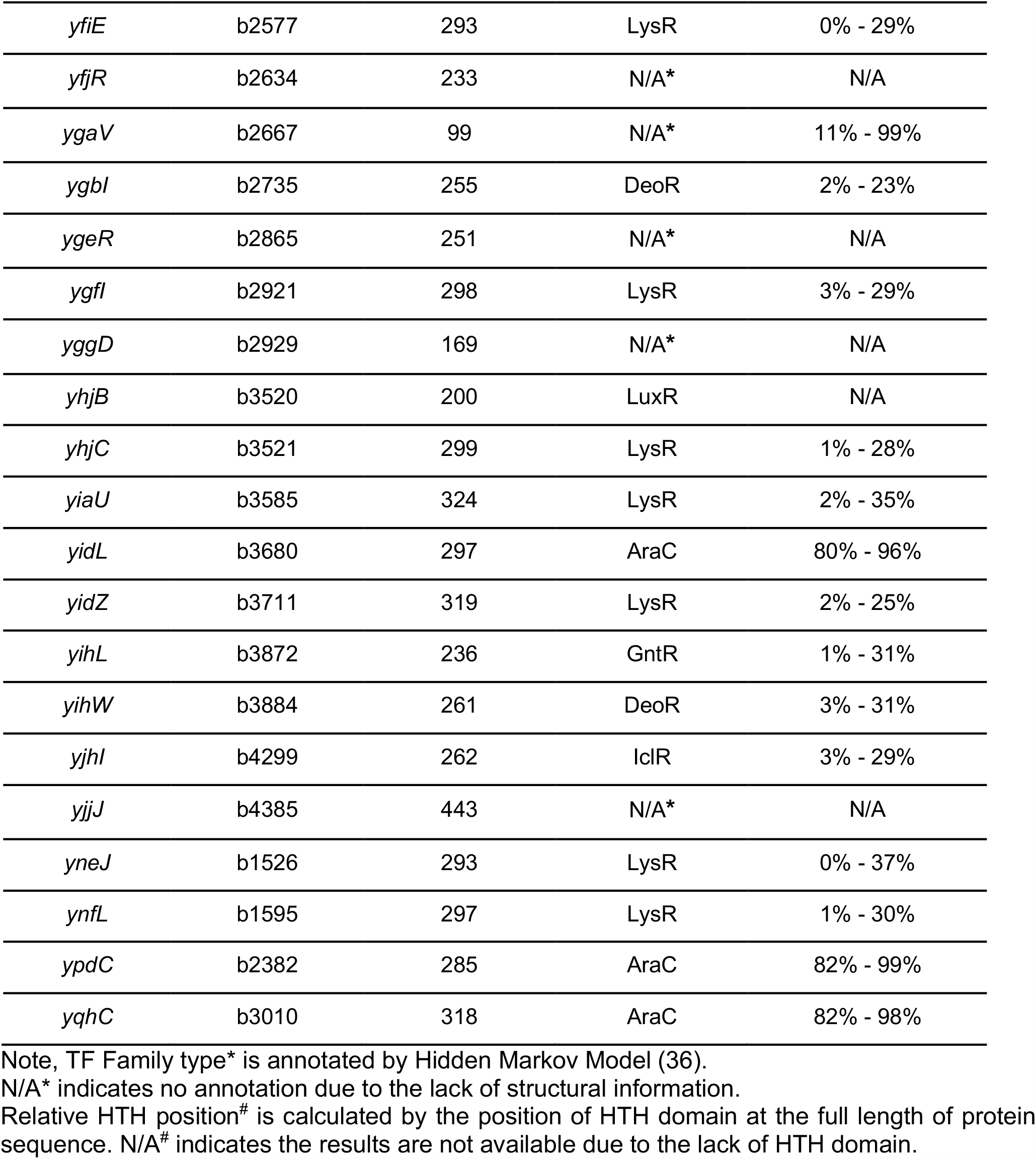
Overview of 40 candidate TFs with the predicted location of the helix-turn-helix (HTH) domain

To predict the putative protein family type of candidate TFs, we employed Hidden Markov Models to annotate them based on the homology to the collection of known protein structures in the SUPERFAMILY 2 database (37) (**Table 1, Dataset S7**). We found that the majority of these 40 candidate TFs contain winged HTH DNA-binding domains, and can be grouped into different TF family types based on homology to known transcription factors (**Supplementary Figure 2A, Supplementary Table 1**) (36). These candidates can be classified into nine known TF family types (LysR, AraC, GntR, CheY, TetR, LuxR, GalR/LacI, IclR, DeoR) and one unknown group (due to the lack of structure information), which were listed in “TF family type” (**Supplementary Figure 2A**). We then calculated the relative position of the helix-turn-helix (HTH) domain for all the candidate TFs, according to the start and end position of HTH domain (amino acids sequence) (11) (**Supplementary Figure 2B**). Several candidate TFs (YfjR, YgeR, YggD, YhjB, YjjJ) do not have a predicted DNA-binding domain due to a lack of structural information, the relative HTH positions of which were annotated as N/A.

### Identification of the genome-wide binding sites for candidate TFs

Having predicted the structural properties of the 40 candidate TFs, we next characterized their genome-wide DNA-binding sites. We constructed 40 myc-tagging strains corresponding to each candidate TF of interest and employed a multiplexed chromatin immunoprecipitation combined with lambda exonuclease digestion (Multiplexed ChIP-exo) to detect the large-scale DNA binding sites under the experimental conditions (**Supplementary Figure 3**) (13).

We examined the genome-wide binding profiles for all candidate TFs using the peak-calling algorithm MACE (17), which confirmed that 34 out of 40 were DNA-binding proteins (**Figure 2A**). A total of 588 reproducible binding sites were identified for verified DNA-binding proteins (**Figure 2B**). Four of the six unconfirmed candidates, YgeR, YggD, YjjJ, and YfjR were annotated as non-HTH domain proteins by Hidden Markov Models (36), which could possibly explain why they did not bind to the genome. Conversely, given the high homology to known DNA-binding proteins, it is possible that YpdC and YeeY are putative TFs but are not active under the experimental conditions used in this study.

**Figure 2.**
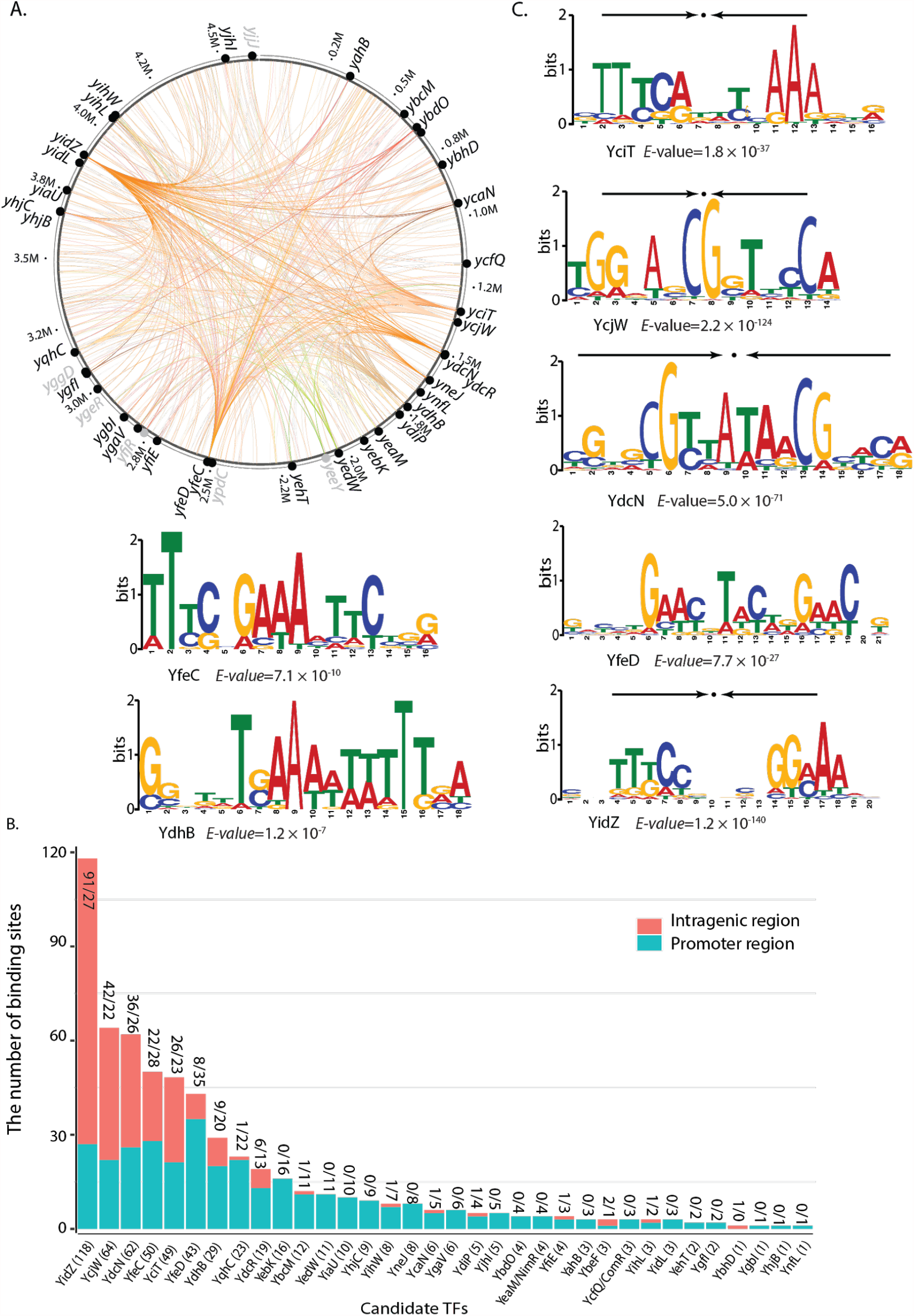
A global landscape of DNA binding events for uncharacterized TFs during growth at active conditions. (A) Binding sites identified by ChIP-exo are mapped onto the *E. coli* genome to provide a network-level perspective of binding activity. Experimentally verified candidate TFs are shown in black, uncharacterized TFs without binding peaks under the growth conditions are shown in grey. Each line indicates the interaction between the TF and the target gene. (B) 34 validated TFs show a varied number of binding sites between the intragenic region and promoter region. The numbers (#/#) above the bar indicate the number of sites that are located at the intragenic region and promoter region, respectively. The number (#) behind the TF name in the x-axis is the total number of binding sites for each validated TF. (C) The consensus sequence motifs for seven uncharacterized TFs determined by ChIP-exo. The height of the letters (in bits on the y-axis) represents the degree of conservation at a given position within the aligned sequence set, with perfect conservation being 2 bits. Arrows above motif indicate the presence of palindromic sequences.

For the 34 validated candidate TFs, we analyzed the conserved binding motifs using the MEME algorithm (38) (**Figure 2C**). We found that the consensus binding motifs for YciT, YcjW, YdcN, and YidZ were palindromic, an observation consistent with the structural predictions that these TFs likely form dimers or tetramers that facilitate binding to specific DNA sequences (**Supplementary Table 2**). Though the remaining validated TFs had a small number of binding peaks, we found similarities of the binding sequences exist within the binding peaks for YbcM, YbdO, YcaN, YcfQ, YdiP, YedW, YihW, and YqhC (**Supplementary Figure 4**).

**Table 2.** The classification of representative candidate TFs and proposed functions in *E. coli* K-12 MG1655

| Gene<sup>#</sup><br>(b-number) | Classification<br>of candidate<br>TFs<br>(# of TFBSs) | Family<br>Type | Binding sites<br>associated with<br>metabolic<br>pathway | Proposed<br>regulatory roles | Results |
| --- | --- | --- | --- | --- | --- |
| *yidZ*<br>(b3711) | Type I<br>(118) | LysR | Widespread,<br>intragenic binding | Target genes have<br>diverse functions | Figure 4 |
| *yfeC*<br>(b2398) | Type II<br>(50) | N/A* | *chaAB, panD, grxC,*<br>*pqqL, hybE, lpp,*<br>*rpmH, rpmB* | *yfeC* mutant was<br>reported to increase<br>eDNA release (43) | Figure 5 |
| *yciT*<br>(b1284) | Type II<br>(49) | DeoR | *ybiO, ybiV, ybiY* | A regulator involved<br>in osmolarity | Figure 6 |
| *ydhB*<br>(b1659) | Type II<br>(29) | LysR | *ydhB, ydhC* | A regulator involved<br>in purine<br>metabolism | Supplementary<br>Figure 5 |
| *ybcM*<br>(b0546) | Type II<br>(12) | AraC | *ybcL, ucpA* | A regulator related<br>to stress response | Figure 7 |
| *yneJ*<sup>#</sup><br>(b1526) | Type II<br>(8) | LysR | *sad, yneJ* | A regulator involved<br>in glutamate<br>metabolism | Supplementary<br>Figure 6 (44) |
| *yjhI*<sup>#</sup><br>(b4299) | Type II<br>(5) | IclR | *yjhG, yjhH, yjhI* | A regulator related<br>to the energy<br>conversion between<br>pyruvate and<br>glycolaldehyde | Supplementary<br>Figure 7 |
| *yfiE*<sup>#</sup><br>(b2577) | Type II<br>(4) | LysR | *yfiE, eamB* | A regulator related<br>to the control of a<br>cysteine and O-acetylserine<br>exporter | Supplementary<br>Figure 8 |
| *ygbI*<br>(b2735) | Type III<br>(1) | DeoR | *ygbJ, ygbK* | A regulator related<br>to four-carbon acid<br>sugar catabolism | Figure 8 |
| *ynfL*<br>(b1595) | Type III<br>(1) | LysR | *ynfL, ynfM* | A regulator involved<br>in the control of<br>arabinose efflux<br>transporter | Supplementary<br>Figure 9 |

\*N/A indicates no prediction due to the lack of structural information. Genes<sup>#</sup> were analyzed and presented in the supplementary material.

### Mapping interactions between RNA polymerase and candidate transcription factors

Gene expression relies on promoter recognition by the RNAP holoenzyme and subsequent transcription initiation. Bacterial promoters consist of at least three RNAP recognition sequences: the − 10 element, the − 35 element, and the UP element (39, 40). The -10 element is optimally positioned between 7 and 12 bp upstream of the transcription start site. The − 10 and − 35 elements are recognized by the RNAP ς subunit (41), and the UP element, located upstream of the − 35 element, is recognized by the RNAP α subunit (39, 42). The RNAP holoenzyme consists of a core enzyme and one of the sigma factors in bacteria. In *E. coli*, the majority of the promoter regions are bound by RNA polymerase sigma factor RpoD (σ^70^) (15). Thus, we performed ChIP-exo experiments to identify genome-wide binding peaks for RNAP and σ^70^ (**Figure 3A**). To examine transcription initiation of the binding sites of candidate TFs, we defined three interactions between RNA polymerase, RpoD, and candidate TFs: (i) RNAP + RpoD: a binding site is located upstream of a target gene, and both RNAP and RpoD recognize the promoter region of this gene; (ii) RNAP_only: a binding site is located upstream of the coding region of a target gene, but only RNAP recognizes the promoter region (while RpoD could not recognize the promoter region, it is likely that alternative sigma factors recognize this promoter region); and (iii) others: includes two interactions; one where a binding site is located within the coding region, the other where a binding site is located within the upstream of the coding region but neither RNAP or RpoD recognize the promoter region. Given these criteria, we identified 208 binding sites belonging to type (i) and 75 binding sites belonging to type (ii). Thus, a total of 283 binding sites overlap with RNAP for the 34 candidate TFs, accounting for 48% (283/588) of total binding sites (**Figure 3B**).

**Figure 3.**
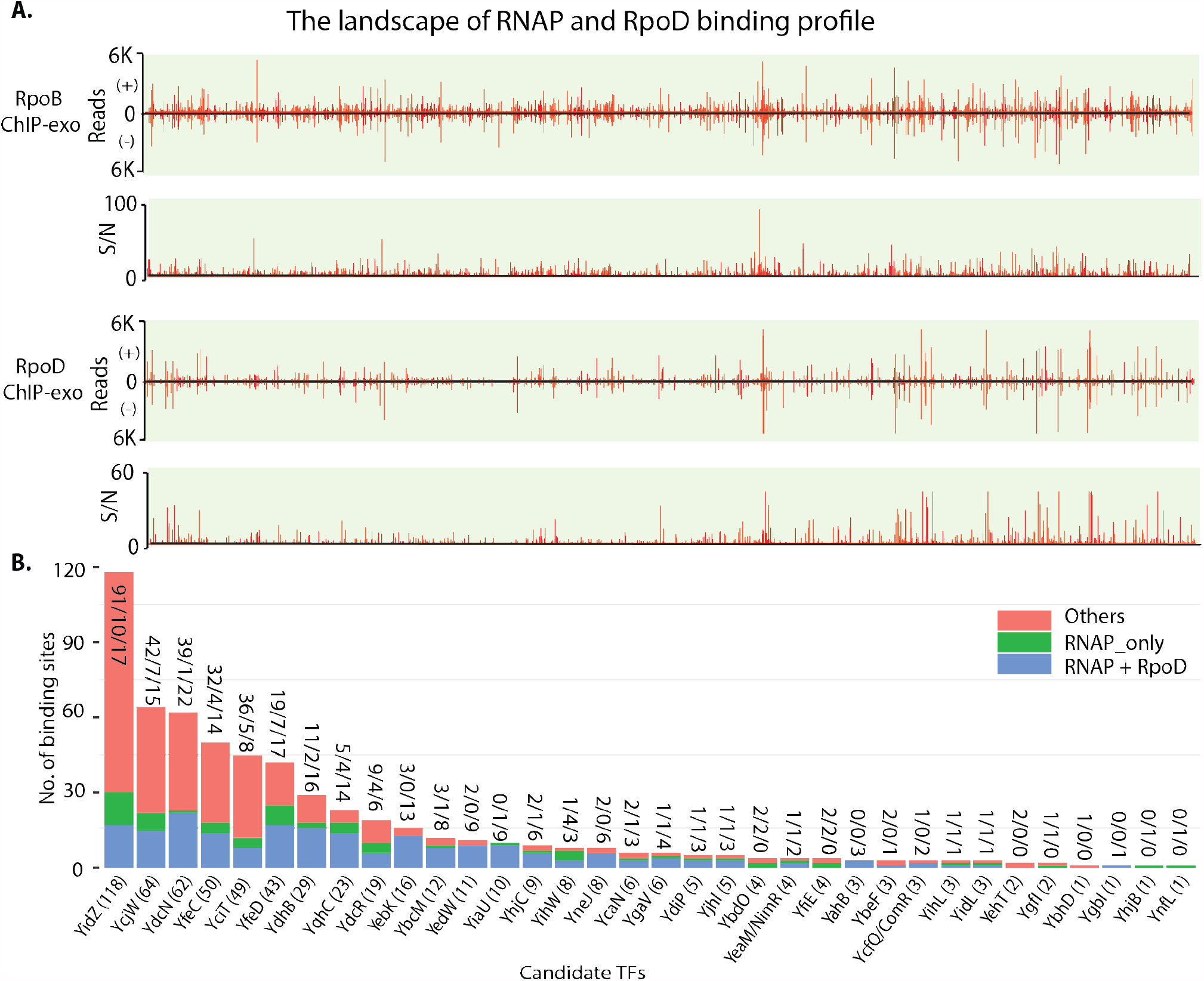
Mapping genome-wide binding sites with RNA polymerase activities. (A) The landscape of RNAP and RpoD binding sites at the genome in *E. coli* K-12 MG1655. Upper panel: overview of RpoB binding sites. Bottom panel: overview of RpoD binding sites. S/N denotes signal-to-noise ratio. (+) and (−) indicate reads mapped on forward and reverse strands, respectively. (B) The classification of the interactions between RpoB, RpoD, and binding sites for 34 candidate TFs. The number (#/#/#) above the bar indicates the number of sites that fit for different types of interactions (in the order of “Others, RNAP_only, and RNAP + RpoD”). Others include two interactions: one where a binding site is located within the coding region, the other where a binding site is located within the upstream of the coding region but neither RNAP or RpoD recognize the promoter region. The number (#) behind the TF name in the x-axis is the total number of binding sites for each validated TF.

### Deciphering regulatory targets of candidate transcription factors

Having verified whether candidate TFs were DNA-binding proteins, we next assessed their putative functions. We used the definition put forth by Shimada et al.—based on the number of target genes—to classify the regulatory nature of the TFs studied here (10). This definition uses four classes: (i) nucleoid-associated regulators (hundreds of target genes); (ii) global regulators (>100 target genes); (iii) local regulators (<100 target genes); and (iv) single-target regulators. Accordingly, we classified the 34 validated TFs into three types of regulators based on the number of target genes identified: global regulator (type I, >100 target genes); local regulator (type II, <100 target gene); and single-target regulator (type III). We inferred the putative biological functions of ten validated TFs based on the genome-wide binding sites (**Table 2**). To confirm the regulatory roles, five of ten validated TFs in the three categories—one global regulator (YidZ), three local regulators (YfeC, YciT, and YbcM), and one single-target regulator (YgbI)—were selected as the representative TFs for detailed analysis. The remaining five validated TFs can be found in the **Supplementary Material**. To infer the regulatory roles, we combined the genome-wide binding sites with gene expression profiling to analyze the most significant enrichment of pathways in which validated TFs are involved.

#### A global regulator (type I), YidZ

We identified 118 genome-wide binding sites (**Figure 4A**). All identified peaks had the expected bimodal shape (45). We used MEME to identify the enriched sequence motif of YidZ binding sites, which is a highly significant enrichment to the consensus DNA sequence (38) (*E*-value = 1.2e^-140^, **Figure 2C, Supplementary Figure 10**). Most of the identified binding peaks (108 out of 118 peaks) contain part of or the full consensus motif, indicating that the majority YidZ-DNA binding sites are high confidence. The consensus binding motif is palindromic, suggesting YidZ is capable of forming a dimeric protein *in vivo*. This finding is consistent with the structural predictions (**Figure 4B, Supplementary Table 2**).

**Figure 4.**
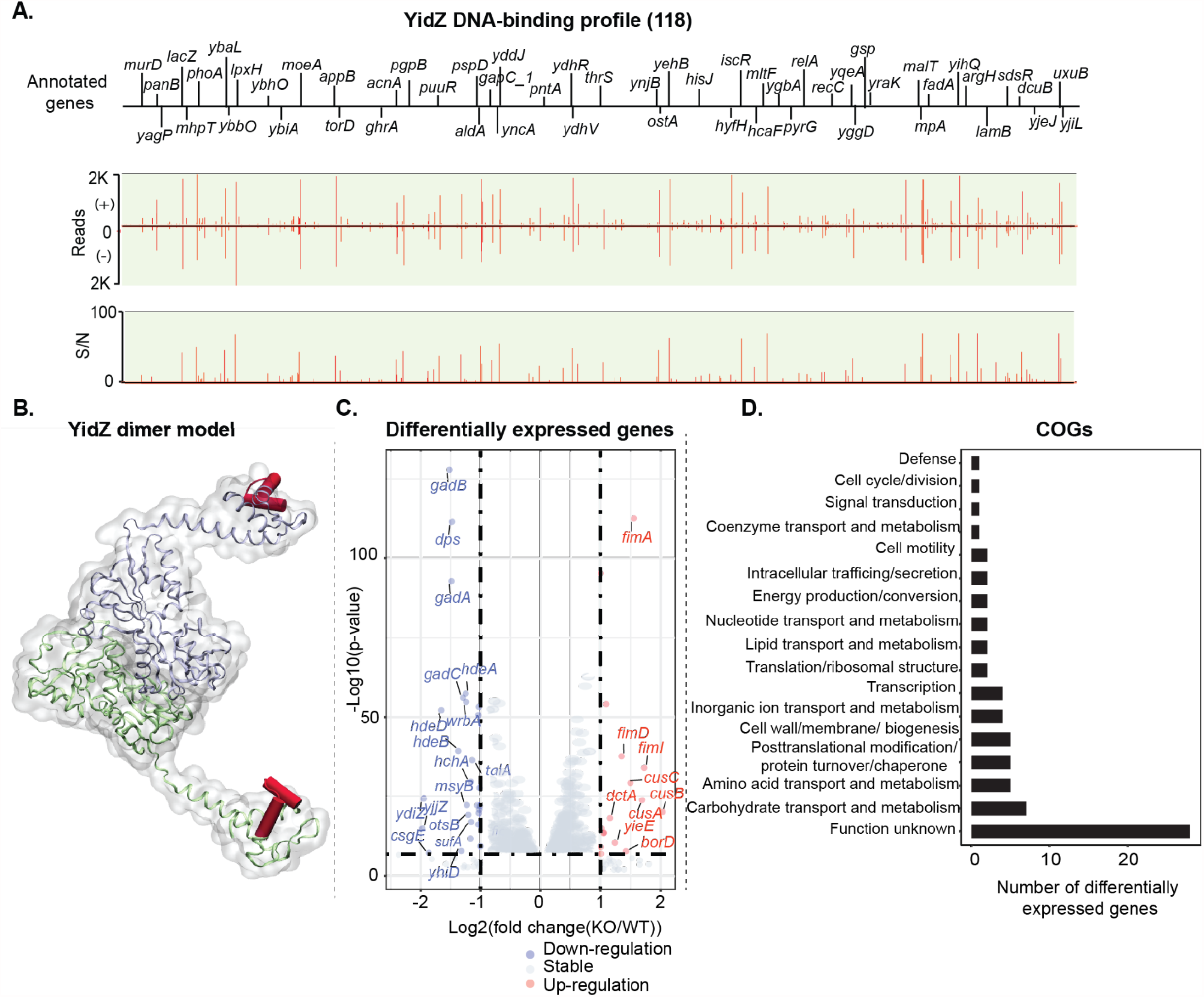
Using YidZ as a representative to illustrate type I global regulator. (A) Genome-wide binding of YidZ identified 118 reproducible binding events at the genome. 77% (91/118) of binding sites are located within the coding region while 23% (27/118) are located within the regulatory region. S/N denotes signal-to-noise ratio. (+) and (−) indicate reads mapped on forward and reverse strands, respectively. (B) Structural analysis of YidZ. YidZ is annotated as a LysR-type putative transcription factor by hidden Markov model. 3D structure is predicted by SWISS-MODEL. The Helix-Turn-Helix domain is highlighted by the red color. (C) 74 genes were differentially expressed after deletion of *yidZ* (cut-off value is log_2_ fold-change >=1, or <=-1, and adjust p-value < 0.05). (D) Functional classification of genes regulated by YidZ. The functions of genes regulated by YidZ are diverse. Additionally, the biological significance of 38% (28/74) of genes is still unknown.

To determine the relative location between YidZ binding *in vivo* and RNA polymerase, we used ChIP-exo to map the genome-wide distribution of RpoB and σ^70^ under the same growth condition used for YidZ. Among 27 intergenic bindings, we found that there are 12 binding sites in the promoters in the presence of core RNAP and σ^70^, 9 binding sites at the promoter in the presence of core RNAP, and 6 binding sites at the promoter DNA in the absence of core RNAP and σ^70^ (**Figure 3A**). Of the 91 intragenic binding sites, 34 are located inside the genes in the presence of core RNAP at the promoter DNA; the remaining 57 binding sites are in the absence of core RNAP at the promoter.

Next, to test whether YidZ impacts the expression of these target genes, we compared the gene expression profile between the wild type strain and the *yidZ* knockout strain using RNA-seq. We noted that 26% (19 of 74) differentially expressed genes were directly regulated by YidZ, though most of the YidZ target genes were not differentially expressed after the deletion of *yidZ* (**Figure 4C**). We also examined the functions of differentially expressed genes and found that operons associated with acid stress and amino acid transport and metabolism (*gadA, gadBC, hdeD, hdeAB-yhiD*) are down-regulated after the deletion of YidZ, while the genes involved in carbohydrate transport and metabolism (*rbsD, malM, malE, malX*) are up-regulated. There are, however, a large number of genes regulated by YidZ with unknown functions (**Figure 4D**).

Overall, we observed two notable features of the YidZ binding profile. First, YidZ has a large number of binding sites, with 77% (91/118) located within the genes and 23% (27/118) located within the intergenic regions. Second, we found that the YidZ binding is associated with diverse gene functions based on Clusters of Orthologous Groups (COGs) annotations (21) (**Figure 4D**). However, we did not find any significantly enriched COGs (p < 0.01), indicating that genes regulated by YidZ are not, as a group, strongly associated with any specific function(s).

#### A local regulator (type II), YfeC

The *yfeC* gene was found by screening the entire Keio collection to identify genes whose deletion increases extracellular DNA (eDNA) production in *E. coli* (43). However, there are no experiments to characterize the interactions between YfeC and DNA. Here, we identified 50 genome-wide binding sites of YfeC in *E. coli* K-12 MG1655 (**Figure 5A**). Functional classification showed that 50 target genes of YfeC are involved in various functional groups, such as nutrient transport and metabolism, DNA replication, transcription, translation, and cell envelope biogenesis (**Figure 5B**). We then enriched the sequence motif of YfeC binding sites (*E*-value=7.1e-10, **Figure 2C**). The consensus DNA binding sequence showed that the TFBS of YfeC enclose TTC-rich inverted repeats separated by 6-nt. It is likely that YfeC can form the homodimer in the cell as inferred from its structure (**Supplementary Figure 11**).

**Figure 5.**
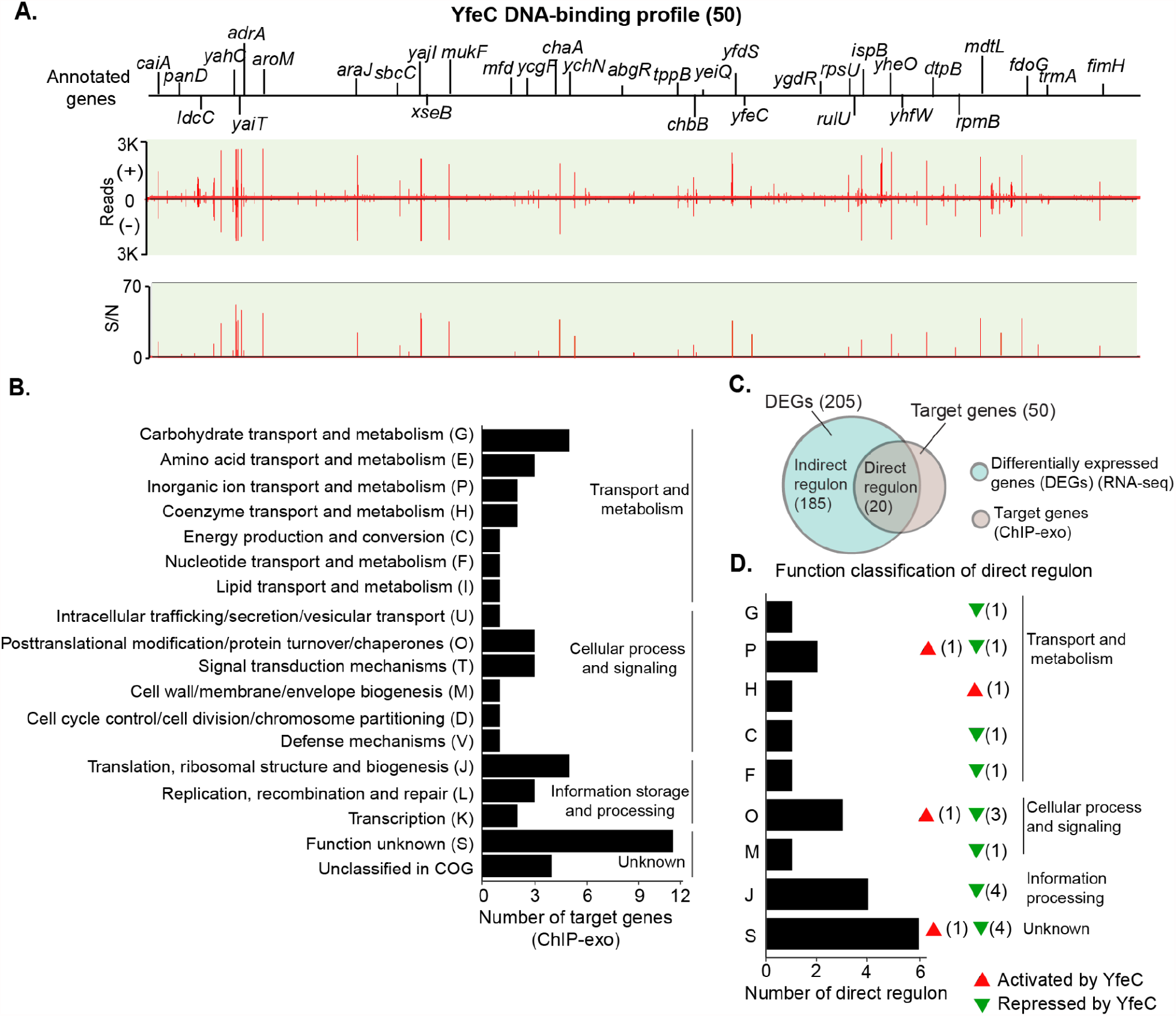
Using YfeC as a representative to illustrate type II regulators. (A) Genome-wide binding of YfeC identified 50 reproducible binding events at the genome. 40% (20/50) of binding sites are located within the coding region while the remaining 60% (30/50) are located within the regulatory region. S/N denotes signal-to-noise ratio. (+) and (−) indicate reads mapped on forward and reverse strands, respectively. (B) Functional classification of target genes from YfeC genome-wide bindings. The enriched functions are in three groups: transport and metabolism pathways, cellular process/signaling, and transcription/translation. (C) Comparison of ChIP-exo results and gene expression profiles to distinguish direct and indirect YfeC regulon under the test condition. (D) Functional classification of genes directly regulated by YfeC. One-letter abbreviations for the functional categories are the same as those in figure 6B. Red triangles represent activation by YfeC. Green triangles represent repression by YfeC. The number behind the triangle represents the number of direct regulon.

To determine a causal relationship between YfeC binding and transcript level, we compared the expression of all genes between wild type and *yfeC* deletion strains. A total of 205 genes were differentially expressed after deletion of *yfeC* (cut-off value is log_2_ fold-change >=1, or <=-1, and adjust p-value < 0.05) **(Figure 5C, Supplementary Figure 12)**. Of 205 differentially expressed genes, 124 (60.2%) of them were up-regulated after the deletion of *yfeC*. Combining YfeC ChIP-exo results with the transcriptome data, we found that 40% (20 of 50) of the genes (or operons) with YfeC binding sites were directly regulated by YfeC under the test conditions (**Figure 5C, Supplementary Table 3**). Of these 20, 80% (16 of 20) of direct regulon are repressed by YfeC, indicating that these genes are up-regulated in the *yfeC* deletion strain (**Figure 5D**). These direct regulon impact various functional groups, such as nutrient transport and metabolism (*chaB, ychO, panD*), translation (*rpmH, rpmB, rpsU*), post-translational modification (*grxC, pqqL, hybE*), and cell envelope (*lpp*). Therefore, the global picture of the YfeC regulon highlights the complexity of the cellular response mediated by this transcriptional regulator. With this information, we demonstrated that the physiological roles of YfeC are those involved in these cellular processes in *E. coli*. The next question is whether any phenotype is caused by the *yfeC* mutant in *E. coli*.

A previous study reported that single-gene deletion strains for genes (*rna, hns, nlpI, rfaD, yfeC*) altered eDNA production in *E. coli*. These mutations were related to general cellular processes, such as transcription (*rna, hns*), lipid transport (*nlpI*), cell envelope (*rfaD*), and unknown function (*yfeC*) (43). These results suggest that the *yfeC* gene is associated with a mutant phenotype-eDNA production in *E. coli*. Furthermore, although the underlying mechanisms remain unknown, the study hints that eDNA release might be related to multiple cellular processes rather than a single biological pathway.

At this point there is no detailed molecular study to determine the mechanism of eDNA release that YfeC regulates in *E. coli*. Designing such a study may serve as the context of future studies.

#### Local regulator (Type II), YciT

YciT was annotated as a DeoR-type putative transcription factor via Hidden Markov Model. However, direct measurement *in vivo* of YciT binding has not been achieved. Therefore, we identified 49 genome-wide binding sites of YciT in *E. coli* K-12 MG1655 (**Figure 6A**), and then enriched the sequence motif of YciT binding sites (*E*-value = 1.8e-37, **Figure 2C**). Similar to the YidZ binding motif, the consensus DNA binding motif is palindromic, suggesting it could form a dimeric protein *in vivo*.

**Figure 6.**
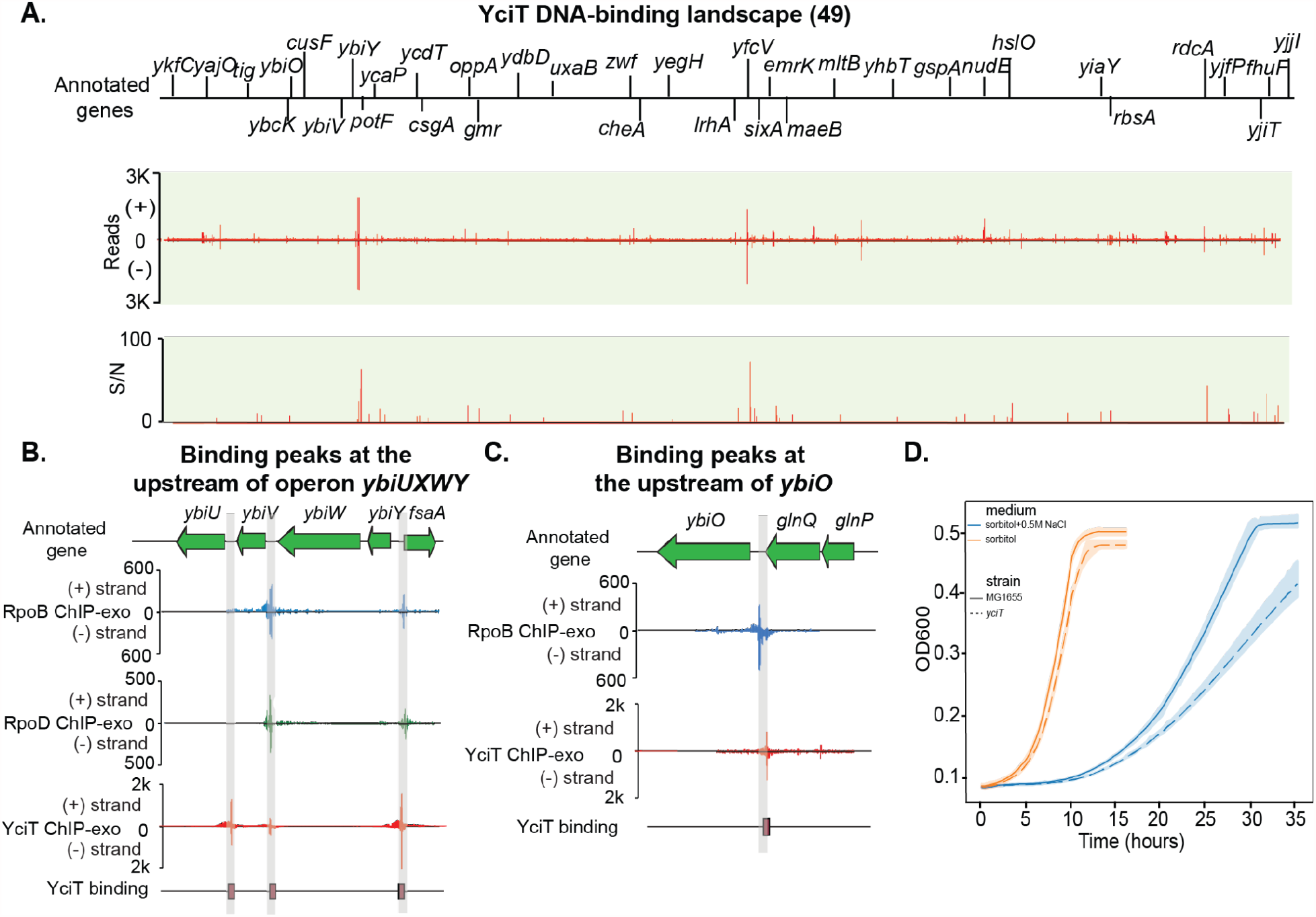
Using YciT as a representative to illustrate type II local regulators. (A) Genome-wide binding of YciT identified 49 binding events at the genome. S/N denotes signal-to-noise ratio. (+) and (−) indicate reads mapped on forward and reverse strands, respectively. (B) YciT binding peak at the upstream of operon *ybiUVWY* and gene *fsaA*. (C) YciT binding peak at the upstream of gene *ybiO*. (D) Growth profiles of the wild type and *yciT* deletion strains in the absence and presence of 0.5 M NaCl in M9 minimal medium with 0.2% (w/v) sorbitol as the sole carbon source. Width of shaded bands represents standard deviation of the corresponding growth trajectory.

To predict the putative functions of YciT, we assessed the genomic location of YciT binding sites and the functions of corresponding target genes using currently available genome annotation. We found 47% (23 out of 49) of binding sites located within regulatory regions, indicating that these binding events may modulate the target genes. The majority of these target genes code for proteins involved in key metabolic pathways -sugar phosphatase (*ybiV*), a putative pyruvate formate-lyase activating enzyme (*ybiY*) and fructose-6-phosphate aldolase1 (*fsaA*) (**Figure 6B**). Some of the other genes encode products involved in membrane components, such as moderate conductance mechanosensitive channel YbiO (*ybiO*), copper/silver export system periplasmic binding protein (*cusF*), and outer membrane protein X (*ompX*) (**Figure 6C**). The remaining genes are unknown functions (*ykfC, ycaP, ydbD*, and *yfdQ*). With this information, we hypothesized that YciT may regulate: (1) target genes (*ybiV, ybiY, fsaA*) involved in the metabolic pathways, and (2) target genes (*ybiO, cusF, ompX*) related to the osmotic stress. Combining the ChIP-exo result with the transcriptome data, we found that these target genes involved in metabolic pathways and membrane components were directly regulated by YciT under the test conditions (**Supplementary Figure 13**), indicating that YciT may participate in the control of the metabolic pathways and (or) osmotic stress in *E. coli*.

To test the hypothesis, we evaluated the impact of *yciT* deletion on the growth of *E. coli* in M9 minimal media containing different carbon sources (glucose, fructose, sorbitol). We found that deletion of the *yciT* gene did not reveal significant growth deficiencies compared to the wild type strain, though the final OD_600_ of the *yciT* deletion strain at the stationary phase was slightly lower than the wild type strain (**Supplementary Figure 14**). This observation suggests that these carbon sources are not natural substrates of enzymes (FsaA, YbiV, YbiY). A previous work reported that fructose-6-phosphate aldolase is related to a novel group of bacterial transaldolases (46). However, the physiological role of FsaA is not yet fully understood. Due to the rare, unidentified enzyme activities and substrate, little is known about the impact of YciT on the metabolic pathways.

We then assessed the effects of osmotic stress on *E. coli* grown in the M9 minimal medium with sorbitol (0.2% w/v) as the sole carbon source (**Figure 6D**). We found osmotic stress induced growth retardation in the wild type and *yciT* deletion strains. Specifically, high osmolarity resulted in impaired growth and slowed the growth rate of the *yciT* deletion strain. Thus we demonstrated that YciT is involved in the control of osmolarity in *E. coli*.

#### Local regulator (type II), YbcM

The *ybcM* gene of *Escherichia coli* was found by screening genes whose products protect *E. coli* from lethal effects of stresses. A previous study reported that the *ybcM* mutant showed hyper lethal phenotype under sodium dodecyl sulfate (SDS) stress, which induces oxidative stress (47). But there are no *in vivo* experiments to confirm its DNA binding capacity. To determine the binding sites, the ChIP-exo experiment for YbcM was conducted under oxidative stress. We then identified 12 genome-wide binding sites in *E. coli* K-12 (**Figure 7A**). 92% (11/12) of the binding sites are located upstream of target genes. We found one binding site located upstream of operon *ybcLM*, indicating autoregulation of *ybcM* (**Figure 7B**). The gene *ybcL* encodes the periplasmic protein YbcL, and has sequence and structural similarity to rat/human RKIP (Raf kinase inhibitor protein), which modulates signal transduction pathways (48).

**Figure 7.**
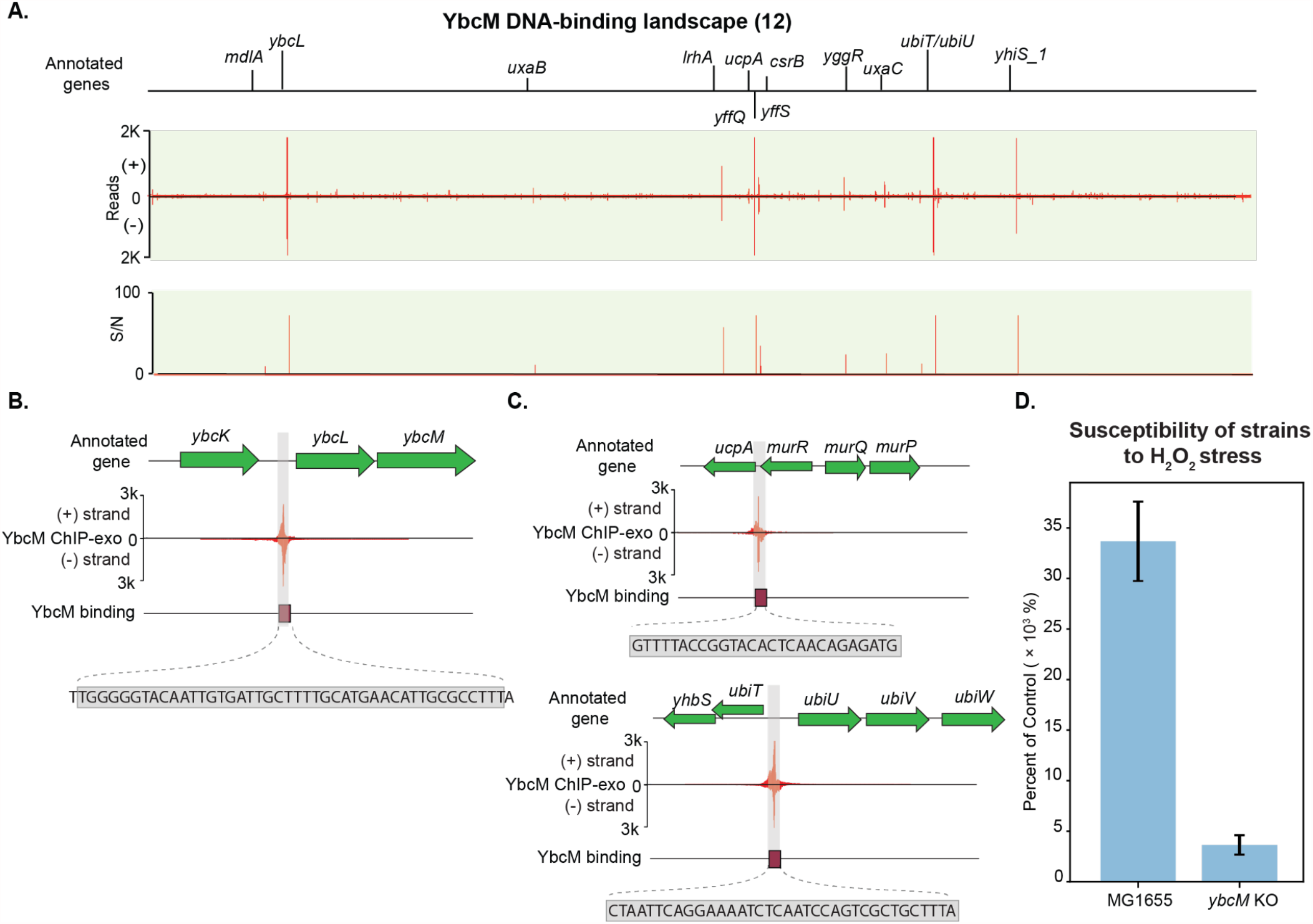
Using YbcM as a representative to illustrate type II regulators. (A) Genome-wide binding of YbcM identified 12 reproducible binding events at the genome. S/N denotes signal-to-noise ratio. (+) and (−) indicate reads mapped on forward and reverse strands, respectively. (B) In-depth mapping of YbcM binding site explains how YbcM interacts with the upstream of operon *ybcLM*. The rectangle denotes the sequence recognized by YbcM. (C) Zoom-in binding peaks of YbcM at the upstream of genes *ucpA* and *ubiT*. (D) Susceptibility of wild type and *ybcM* deletion strains under the oxidative stress. Both wild type and *ybcM* KO strains (mid-log phase cells) were treated with 60 mM H_2_O_2_ for 15 min. The sensitivity of cells to the lethal effects was expressed as percent survival of treated cells relative to that of untreated cells determined at the time of treatment. Wild type strain showed 7 times higher survival rate than that of *ybcM* KO strain.

To predict the functions of YbcM, we examined 12 binding sites and their functions. We found that there are two important binding sites involved in stress response. The first was located upstream of the gene *ucpA*, encoding the oxidoreductase UcpA. YbcM was found to bind the upstream of *ucpA* (**Figure 7C, upper panel**). Overexpression of *ucpA* in plasmids was previously shown to lead to improved tolerance to furan (49), a chemical likely generating oxidative stress. The other divergent binding site was located between operons *ubiT-yhbS* and *ubiUV* (**Figure 7C, bottom panel**). Here, the *ubiT* gene encodes anaerobic ubiquinone biosynthesis accessory factor UbiT, *yhbS* encodes putative N-acetyltransferase YhbS, and *ubiUV* encodes ubiquinone biosynthesis complex UbiUV. Another gene, *ubiW*, nearby operon *ubiUV*, encodes putative luciferase-like monooxygenase. We then identified a consensus YbcM binding motif in the promoter region of these target genes (**Supplementary Figure 4**). Taken together, these target genes suggest that YbcM might be a regulator involved in the oxidative stress in *E. coli*.

To confirm the physiological role, the survival rate of the wild type and *ybcM* deletion strains were compared under oxidative stress conditions (**Figure 7D**). The survival rate of the wild strain was 8-fold higher over the *ybcM* deletion strain after 15 min 60 mM H_2_O_2_ treatment. This observation supports the involvement of YbcM in the reactive oxygen species (ROS) stress response. Therefore, this data supports that YbcM is involved in the oxidative stress response.

#### A single-target regulator (type III), YgbI

YgbI was annotated as a putative transcription factor via Hidden Markov Model. However, *in vivo* analysis of direct interactions between YgbI and DNA in *E. coli* has not been reported. Here, we identified a single divergent binding site between the *ygbI* and *ygbJ* genes, indicating autoregulation of *ygbI* (**Figure 8A**). To explore how the YgbI binding site regulates target genes, we compared the interaction between the YgbI binding site and RNAP binding site at the promoter region, and found that YgbI binding overlaps the promoter region recognized by RNAP. This suggested that YgbI could block RNAP binding to the promoter or inhibit the transcription initiation, repressing the expression of downstream genes (*ygbJ, ygbK*).

**Figure 8.**
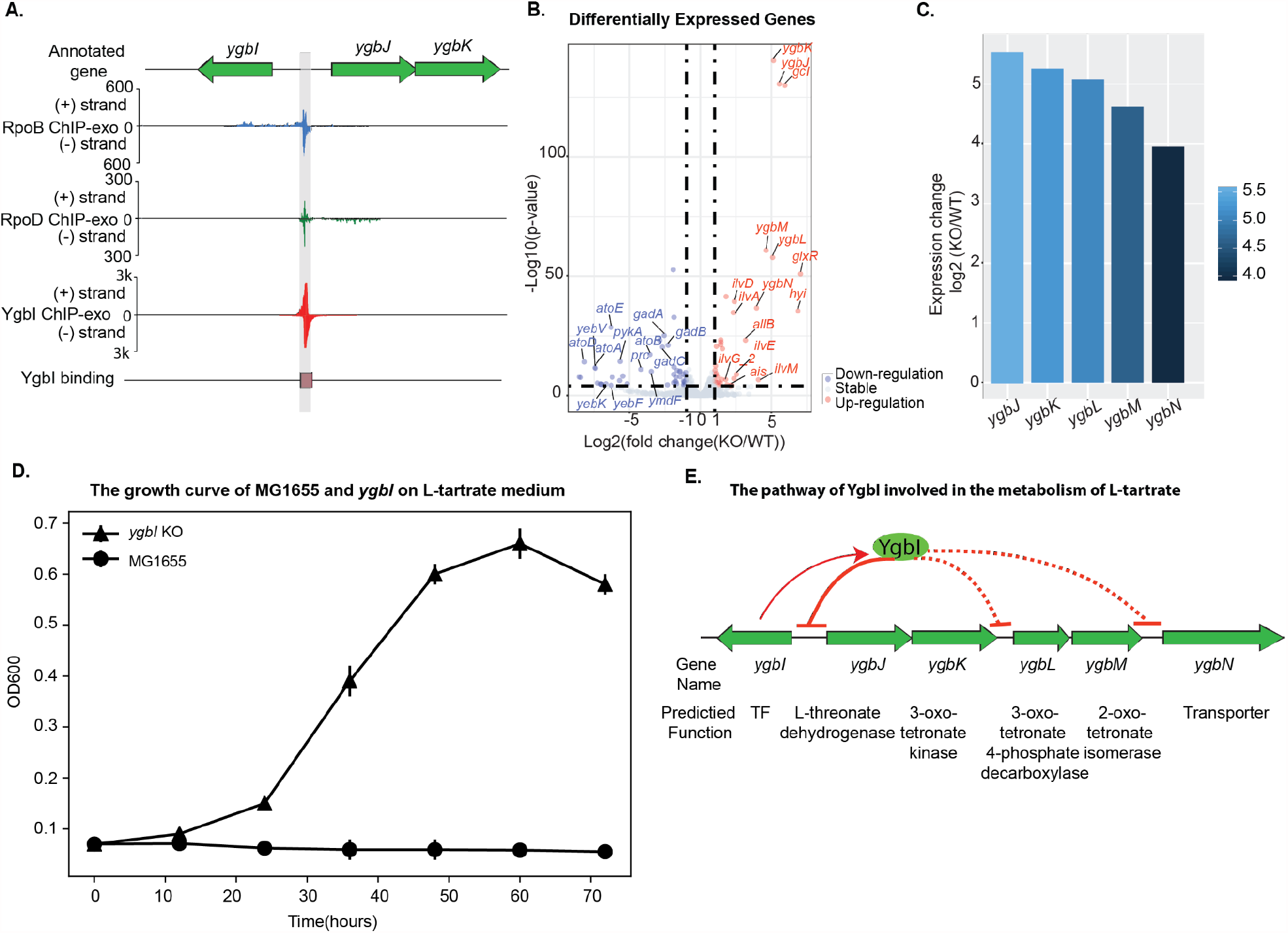
Using YgbI as a representative to illustrate type III regulators (single-target). (A) Zoom-in mapping of YgbI, RpoB, and RpoD binding events explains how YgbI binds onto the upstream of operon *ygbJK*, covering the region that RpoB can recognize. Thus YgbI blocks the transcription initiation of operon *ygbJK* when YgbI is active. (B) 113 genes were differentially expressed after deletion of *ygbI* (cut-off value is log_2_ fold-change >=1, or <=-1, and adjust p-value < 0.05). (C) Expression changes for genes in the *ygbI* deletion strain in a set of genes (*ygbJKLMN*) near the binding peak, compared to the wild type strain. (D) Growth of *E. coli* K-12 MG1655 and *ygbI* deletion strains on 20 mM L-tartrate medium, a four-carbon sugar. Circle markers represent growth of the wild type strain. Triangle markers represent growth of the *ygbI* deletion strain. (E) The proposed mechanism for the regulatory role of YgbI. When YgbI is present (active), it directly represses the promoter of the operon *ygbJK*, and indirectly inhibits the expression of operon *ygbLM* and gene *ygbN*. When gene *ybgI* is knocked out, it leads to de-repression of the operons *ygbJK, ygbLM* and *ygbN*.

To examine this assumption about the regulation of YgbI, we compared gene expression profiling between the wild type and the *ygbI* mutant (**Figure 8B**). The results showed that the expression of a cluster of genes (*ygbJ, ygbK, ygbL, ygbM, ybgN*) are upregulated after the deletion of *ygbI* (**Figure 8C**). These data suggested that YgbI regulates the downstream gene cluster (*ygbJKLMN*) as a repressor, which is consistent with the prediction of a regulatory effect.

Previous studies reported that the downstream gene cluster (*ygbJKLM*) had putative functions for four-carbon sugar catabolism (50), and hypothesized that *E. coli* K-12 strain carrying mutations in the *ygbI* gene would provide a growth benefit on the tartrate medium (51). To verify the function of YgbI, the growth profiles of the wild type and the *ygbI* deletion strain were measured in 20 mM L-tartrate medium. The data showed that the deletion of gene *ygbI* demonstrated the capability for growth on L-tartrate. However, the wild type strain does not grow on L-tartrate medium (**Figure 8D**). Taking these factors into consideration, the potential pathway that YgbI is involved in was proposed as follows: when YgbI is present and active *in vivo*, it directly binds to the promoter of the operon *ygbJK*, and indirectly inhibits the expression of the genes *ygbLM* and *ygbN*. When the gene *ybgI* is knocked out, it leads to de-repression of operons *ygbJK* and *ygbLM* and the gene *ygbN* (**Figure 8E**). Based on the putative function of genes (*ygbJKLMN*), we demonstrated that YgbI is a transcriptional repressor involved in catabolic pathways for four carbon acid sugars in *E. coli* K-12 MG1655.

## Discussion

Despite extensive research over many decades focused on the *E. coli* genome, around 35% of its genes are still poorly characterized, including some uncharacterized transcription factors (9, 52). Our primary goal was to generate a large data set to further identify DNA-binding proteins from a pool of uncharacterized proteins in *E. coli*. In this study, we used a systematic approach to validate 34 computationally predicted transcription factors in *E. coli* K-12 MG1655. We employed a multiplexed ChIP-exo method to characterize genome-wide binding sites and classify this experimental evidence for each TF. Next, we compared the binding profiles of the candidate TFs with binding peaks for RNAP holoenzyme, which generate a total of 283 out of 588 sites that are likely to be regulating a close-by promoter (**Dataset S4**), and provide a coarse-grained functional prediction. Finally, we inferred the putative functions for ten of these candidate TFs (YidZ, YfeC, YciT, YdhB, YbcM, YneJ, YjhI, YfiE, YgbI, YnfL), and verified the biological roles of the representative TFs with detailed analysis. The implications of our results are below.

First, our study collected a large dataset of 588 TFBS, and expanded the total number of verified TFs in *E. coli* K-12 MG1655, close to the estimated total number of 280 (**Supplementary Figure 15**). Comparative analysis of genome-wide binding sites of the TFs and RNAP enables the identification of target genes that are recognized by RNA polymerase complexes. Of the total 588 TFBS, 283 RNAP binding sites represent that almost half of binding sites are likely to be regulating a close-by promoter under the test conditions. Also, the interaction between RNAP and recognition sequence at the promoter region may change depending upon the test conditions. It is possible that some TFBSs that are not identified by RNAP may be recognized by the RNAP complex under different conditions. Further, discovering all of the TFs is fundamental to fully understand the key role TRNs play in enabling bacteria to modulate the expression of thousands of genes in response to environmental and genetic perturbations (6). This study has brought us closer to revealing the identity of all the TFs in *E. coli* K-12 MG1655.

Second, we used the definition of TFs reported by Shimada et al., to classify candidate TFs into three groups: type I regulators, type II regulators, and type III single-target regulators (10). This classification was based on the number of genes regulated by TFs as determined from the systematic evolution of ligands with exponential enrichment (SELEX) (56). Our rationale for using this classification was twofold: (i) the multiplexed ChIP-exo method employed here offers a similar readout to SELEX (i.e., the number of target genes), allowing for its application in the same context; and (ii) it has a successful track record of assigning annotations (e.g., “global” or “local” regulator) prior to a full understanding of the functions of the validated TFs, helping to guide their future study. Thus, we employed this classification based on the number of target genes shown by genome-wide experiments. We expect that a detailed characterization of these validated TFs will help us develop a comprehensive understanding of transcriptional regulation in *E. coli*.

Third, we did not identify binding sites for six of the candidate TFs tested in this study (YgeR, YggD, YjjJ, YfjR, YeeY, YpdC). This result may be due to two reasons. The first is the false-positive predictions of candidate TFs due to the limitations of the sequence homology search. Since the computational predictions were made, studies focusing on the function of four of these six candidates have appeared. YgeR has been recently re-annotated as putative lipoprotein involved in septation (54). YggD has been verified as fumarase E (57). Overexpression of YjjJ increases toxic effects in *E. coli*, thus *yjjJ* is likely to be a toxin (58). YfjR is predicted as a putative TF involved in biofilm formation (59), but a recent study that searched novel TFs involved in biofilm formation has not validated this prediction (32). A second reason for failed prediction is that we may need to test for DNA-binding activity under the right activation conditions. YeeY and YpdC are annotated as a LysR-type regulator with a C-terminal HTH domain and an AraC-type regulator with a C-terminal HTH domain, respectively (**Table 1**), and thus they may have regulatory functions under the appropriate growth conditions.

Fourth, while we identified additional TFs with our experimental data, we did not fully decipher mutant phenotypes. For example, we identified YciT as a TF and found that it directly regulated multiple target genes (*fsaA, ybiY, ybiV*). This result hinted at an uncharacterized pathway composed of genes encoding DUF1479 domain-containing protein (*ybiU*), a sugar phosphatase (*ybiV*), a putative pyruvate formate lyase (PFL) (*ybiW*), a putative pyruvate formate-lyase activating enzyme (PFL-AE) (*ybiY*), and a fructose-6-phosphate aldolase1 (FSA) (*fsaA*) (**Supplementary Figure 16**). However, these enzymes and their corresponding substrates are rare and have not been identified, little is known about their physiological roles in *E. coli* (46). These bottlenecks may pose challenges in fully examining mutant phenotypes. Studying these enzymes should provide insight into biological roles of YciT.

Finally, a collection of data sets about TFBS will lay the foundation for understanding the mechanisms of transcription regulation. In this study, we found YfeC regulates multiple cellular processes in *E. coli*. Previous studies had not dived into a possible relation between eDNA and YfeC. So, we employed a *yfeC* mutant to better understand any possible connections. Consider the common mechanism of eDNA release in bacteria - this through membrane vesicles (MVs) secretion (43). eDNA production relies on several biological processes: (1) DNA replication, to produce DNA for the secretion (refer as to eDNA); (2) nutrient transport and metabolisms, to generate lipid metabolisms for MVs; (3) energy conversion, to produce the energy for conversion of metabolisms and secretion of MVs; (4) transcription and translation, to produce the proteins for the assembly of MVs; (5) post-translational modification, protein turnover, and chaperones, to modify and fold the proteins for the secretion; and (6) cell wall/envelope biogenesis, to recover the cell wall after the secretion of eDNA (**Supplementary Figure 17**) (60). As a repressor, YfeC participates in multiple cellular processes, including lipid metabolism, translation, post-translational modification, and cell wall/envelope biogenesis. Accordingly, these corresponding biological processes are up-regulated after the deletion of *yfeC*. We proposed that the deletion of the *yfeC* gene may speed up cellular processes, leading to eDNA release in *E. coli*.

This study focused on the remaining uncharacterized TFs in *E. coli* K-12 MG1655. Taken together, it significantly expands the size of the potential TFs with experimental evidence, broadening our knowledge of transcriptional regulation in *E. coli*.

## Supplementary materials

See the supplementary materials

## Data availability

The whole dataset of ChIP-exo and RNA-seq has been deposited to GEO with the accession number of GSE159777 (secure token: klinmsskfzgbtwd) and GSE159658 (secure token: kvydaewevbgxtmp), respectively.

## Acknowledgements

We thank Richard Szubin for help with ChIP-exo and RNA-seq library sequencing, Drs. Donghyuk Kim, Sang Woo Seo and Byung-Kwan Cho for the insights, and Marc Abrams for reviewing and editing the manuscript.

## References

1. Browning, D.F. and Busby, S.J.W. (2016) Local and global regulation of transcription initiation in bacteria. Nat. Rev. Microbiol., 14, 638–650.

2. Mejía-Almonte, C., Busby, S.J.W., Wade, J.T., van Helden, J., Arkin, A.P., Stormo, G.D., Eilbeck, K., Palsson, B.O., Galagan, J.E. and Collado-Vides, J. (2020) Redefining fundamental concepts of transcription initiation in bacteria. Nat. Rev. Genet., 10.1038/s41576-020-0254-8.

3. Balleza, E., López-Bojorquez, L.N., Martínez-Antonio, A., Resendis-Antonio, O., Lozada-Chávez, I., Balderas-Martínez, Y.I., Encarnación, S. and Collado-Vides, J. (2009) Regulation by transcription factors in bacteria: beyond description. FEMS Microbiol. Rev., 33, 133–151.

4. Park, P.J. (2009) ChIP–seq: advantages and challenges of a maturing technology. Nature Reviews Genetics, 10, 669–680.

5. Rhee, H.S. and Pugh, B.F. (2012) ChIP-exo method for identifying genomic location of DNA-binding proteins with near-single-nucleotide accuracy. Curr. Protoc. Mol. Biol., Chapter 21, Unit 21.24.

6. Martínez-Antonio, A., Janga, S.C. and Thieffry, D. (2008) Functional organisation of Escherichia coli transcriptional regulatory network. J. Mol. Biol., 381, 238–247.

7. Dobrin, R., Beg, Q.K., Barabási, A.-L. and Oltvai, Z.N. (2004) Aggregation of topological motifs in the Escherichia coli transcriptional regulatory network. BMC Bioinformatics, 5, 10.

8. Fang, X., Sastry, A., Mih, N., Kim, D., Tan, J., Yurkovich, J.T., Lloyd, C.J., Gao, Y., Yang, L. and Palsson, B.O. (2017) Global transcriptional regulatory network for Escherichia coli robustly connects gene expression to transcription factor activities. Proc. Natl. Acad. Sci. U. S. A., 114, 10286–10291.

9. Gao, Y., Yurkovich, J.T., Seo, S.W., Kabimoldayev, I., Dräger, A., Chen, K., Sastry, A.V., Fang, X., Mih, N., Yang, L., et al. (2018) Systematic discovery of uncharacterized transcription factors in Escherichia coli K-12 MG1655. Nucleic Acids Res., 46, 10682–10696.

10. Shimada, T., Ogasawara, H. and Ishihama, A. (2018) Single-target regulators form a minor group of transcription factors in Escherichia coli K-12. Nucleic Acids Res., 10.1093/nar/gky138.

11. Pérez-Rueda, E. and Collado-Vides, J. (2000) The repertoire of DNA-binding transcriptional regulators in Escherichia coli K-12. Nucleic Acids Res., 28, 1838–1847.

12. Eichner, J., Topf, F., Dräger, A., Wrzodek, C., Wanke, D. and Zell, A. (2013) TFpredict and SABINE: sequence-based prediction of structural and functional characteristics of transcription factors. PLoS One, 8, e82238.

13. Cho, B.-K., Knight, E.M. and Palsson, B.O. (2006) PCR-based tandem epitope tagging system for Escherichia coli genome engineering. Biotechniques, 40, 67–72.

14. Seo, S.W., Kim, D., Latif, H., O’Brien, E.J., Szubin, R. and Palsson, B.O. (2014) Deciphering Fur transcriptional regulatory network highlights its complex role beyond iron metabolism in Escherichia coli. Nat. Commun., 5, 4910.

15. Cho, B.-K., Kim, D., Knight, E.M., Zengler, K. and Palsson, B.O. (2014) Genome-scale reconstruction of the sigma factor network in Escherichia coli: topology and functional states. BMC Biol., 12, 4.

16. Langmead, B., Trapnell, C., Pop, M. and Salzberg, S.L. (2009) Ultrafast and memoryefficient alignment of short DNA sequences to the human genome. Genome Biol., 10, R25.

17. Wang, L., Chen, J., Wang, C., Uusküla-Reimand, L., Chen, K., Medina-Rivera, A., Young, E.J., Zimmermann, M.T., Yan, H., Sun, Z., et al. (2014) MACE: model based analysis of ChIP-exo. Nucleic Acids Res., 42, e156.

18. Seo, S.W., Kim, D., Szubin, R. and Palsson, B.O. (2015) Genome-wide Reconstruction of OxyR and SoxRS Transcriptional Regulatory Networks under Oxidative Stress in Escherichia coli K-12 MG1655. Cell Rep., 12, 1289–1299.

19. Ogasawara, H., Ohe, S. and Ishihama, A. (2015) Role of transcription factor NimR (YeaM) in sensitivity control of Escherichia coli to 2-nitroimidazole. FEMS Microbiol. Lett., 362, 1–8.

20. Bailey, T.L., Boden, M., Buske, F.A., Frith, M., Grant, C.E., Clementi, L., Ren, J., Li, W.W. and Noble, W.S. (2009) MEME SUITE: tools for motif discovery and searching. Nucleic Acids Res., 37, W202–8.

21. Tatusov, R.L., Galperin, M.Y., Natale, D.A. and Koonin, E.V. (2000) The COG database: a tool for genome-scale analysis of protein functions and evolution. Nucleic Acids Res., 28, 33–36.

22. Ross, M.G., Russ, C., Costello, M., Hollinger, A., Lennon, N.J., Hegarty, R., Nusbaum, C. and Jaffe, D.B. (2013) Characterizing and measuring bias in sequence data. Genome Biol., 14, R51.

23. Quail, M.A., Otto, T.D., Gu, Y., Harris, S.R., Skelly, T.F., McQuillan, J.A., Swerdlow, H.P. and Oyola, S.O. (2011) Optimal enzymes for amplifying sequencing libraries. Nat. Methods, 9, 10.

24. Lawrence, M., Huber, W., Pagès, H., Aboyoun, P., Carlson, M., Gentleman, R., Morgan, M.T. and Carey, V.J. (2013) Software for computing and annotating genomic ranges. PLoS Comput. Biol., 9, e1003118.

25. Love, M.I., Huber, W. and Anders, S. (2014) Moderated estimation of fold change and dispersion for RNA-seq data with DESeq2. Genome Biol., 15, 550.

26. Biasini, M., Bienert, S., Waterhouse, A., Arnold, K., Studer, G., Schmidt, T., Kiefer, F., Gallo Cassarino, T., Bertoni, M., Bordoli, L., et al. (2014) SWISS-MODEL: modelling protein tertiary and quaternary structure using evolutionary information. Nucleic Acids Res., 42, W252–8.

27. The UniProt Consortium (2017) UniProt: the universal protein knowledgebase. Nucleic Acids Res., 45, D158–D169.

28. Humphrey, W., Dalke, A. and Schulten, K. (1996) VMD: visual molecular dynamics. J. Mol. Graph., 14, 33–8, 27–8.

29. Mermod, M., Magnani, D., Solioz, M. and Stoyanov, J.V. (2012) The copper-inducible ComR (YcfQ) repressor regulates expression of ComC (YcfR), which affects copper permeability of the outer membrane of Escherichia coli. Biometals, 25, 33–43.

30. Luhachack, L., Rasouly, A., Shamovsky, I. and Nudler, E. (2019) Transcription factor YcjW controls the emergency H2S production in E. coli. Nature Communications, 10.

31. Yamamoto, K., Nakano, M. and Ishihama, A. (2015) Regulatory role of transcription factor SutR (YdcN) in sulfur utilization in Escherichia coli. Microbiology, 161, 99–111.

32. Ogasawara, H., Ishizuka, T., Hotta, S., Aoki, M., Shimada, T. and Ishihama, A. (2020) Novel regulators of the csgD gene encoding the master regulator of biofilm formation in Escherichia coli K-12. Microbiology, 166, 880–890.

33. Shimada, T., Yamamoto, K., Nakano, M., Watanabe, H., Schleheck, D. and Ishihama, A. (2019) Regulatory role of CsqR (YihW) in transcription of the genes for catabolism of the anionic sugar sulfoquinovose (SQ) in Escherichia coli K-12. Microbiology, 165, 78–89.

34. Kaznadzey, A., Shelyakin, P., Belousova, E., Eremina, A., Shvyreva, U., Bykova, D., Emelianenko, V., Korosteleva, A., Tutukina, M. and Gelfand, M.S. (2018) The genes of the sulphoquinovose catabolism in Escherichia coli are also associated with a previously unknown pathway of lactose degradation. Sci. Rep., 8, 3177.

35. Turner, P.C., Miller, E.N., Jarboe, L.R., Baggett, C.L., Shanmugam, K.T. and Ingram, L.O. (2011) YqhC regulates transcription of the adjacent Escherichia coli genes yqhD and dkgA that are involved in furfural tolerance. J. Ind. Microbiol. Biotechnol., 38, 431–439.

36. Pandurangan, A.P., Stahlhacke, J., Oates, M.E., Smithers, B. and Gough, J. (2019) The SUPERFAMILY 2.0 database: a significant proteome update and a new webserver. Nucleic Acids Research, 47, D490–D494.

37. Gough, J., Karplus, K., Hughey, R. and Chothia, C. (2001) Assignment of homology to genome sequences using a library of hidden Markov models that represent all proteins of known structure. J. Mol. Biol., 313, 903–919.

38. Bailey, T.L., Boden, M., Buske, F.A., Frith, M., Grant, C.E., Clementi, L., Ren, J., Li, W.W. and Noble, W.S. (2009) MEME SUITE: tools for motif discovery and searching. Nucleic Acids Research, 37, W202–W208.

39. Ross, W., Gosink, K., Salomon, J., Igarashi, K., Zou, C., Ishihama, A., Severinov, K. and Gourse, R. (1993) A third recognition element in bacterial promoters: DNA binding by the alpha subunit of RNA polymerase. Science, 262, 1407–1413.

40. Hawley, D.K. and McClure, W.R. (1983) Compilation and analysis of Escherichia coli promoter DNA sequences. Nucleic Acids Res., 11, 2237–2255.

41. Dombroski, A.J., Walter, W.A., Record, M.T., Jr, Siegele, D.A. and Gross, C.A. (1992) Polypeptides containing highly conserved regions of transcription initiation factor sigma 70 exhibit specificity of binding to promoter DNA. Cell, 70, 501–512.

42. Blatter, E.E., Ross, W., Tang, H., Gourse, R.L. and Ebright, R.H. (1994) Domain organization of RNA polymerase alpha subunit: C-terminal 85 amino acids constitute a domain capable of dimerization and DNA binding. Cell, 78, 889–896.

43. Sanchez-Torres, V., Maeda, T. and Wood, T.K. (2010) Global regulator H-NS and lipoprotein NlpI influence production of extracellular DNA in Escherichia coli. Biochemical and Biophysical Research Communications, 401, 197–202.

44. Rodionova, I.A., Gao, Y., Sastry, A., Monk, J., Wong, N., Szubin, R., Lim, H., Zhang, Z., Saier, M.H. and Palsson, B. PtrR (YneJ) is a novel E. coli transcription factor regulating the putrescine stress response and glutamate utilization. 10.1101/2020.04.27.065417.

45. Kharchenko, P.V., Tolstorukov, M.Y. and Park, P.J. (2008) Design and analysis of ChIP-seq experiments for DNA-binding proteins. Nature Biotechnology, 26, 1351–1359.

46. Schürmann, M. and Sprenger, G.A. (2001) Fructose-6-phosphate Aldolase Is a Novel Class I Aldolase from Escherichia coli and Is Related to a Novel Group of Bacterial Transaldolases. Journal of Biological Chemistry, 276, 11055–11061.

47. Han, X., Dorsey-Oresto, A., Malik, M., Wang, J.-Y., Drlica, K., Zhao, X. and Lu, T. (2010) Escherichia coli genes that reduce the lethal effects of stress. BMC Microbiol., 10, 35.

48. Serre, L., Pereira de Jesus, K., Zelwer, C., Bureaud, N., Schoentgen, F. and Bénédetti, H. (2001) Crystal structures of YBHB and YBCL from Escherichia coli, two bacterial homologues to a Raf kinase inhibitor protein. J. Mol. Biol., 310, 617–634.

49. Wang, X., Miller, E.N., Yomano, L.P., Shanmugam, K.T. and Ingram, L.O. (2012) Increased furan tolerance in Escherichia coli due to a cryptic ucpA gene. Appl. Environ. Microbiol., 78, 2452–2455.

50. Zhang, X., Carter, M.S., Vetting, M.W., San Francisco, B., Zhao, S., Al-Obaidi, N.F., Solbiati, J.O., Thiaville, J.J., de Crécy-Lagard, V., Jacobson, M.P., et al. (2016) Assignment of function to a domain of unknown function: DUF1537 is a new kinase family in catabolic pathways for acid sugars. Proc. Natl. Acad. Sci. U. S. A., 113, E4161–9.

51. Guzmán, G.I., Sandberg, T.E., LaCroix, R.A., Nyerges, Á., Papp, H., de Raad, M., King, Z.A., Hefner, Y., Northen, T.R., Notebaart, R.A., et al. (2019) Enzyme promiscuity shapes adaptation to novel growth substrates. Mol. Syst. Biol., 15, e8462.

52. Ghatak, S., King, Z.A., Sastry, A. and Palsson, B.O. (2019) The y-ome defines the 35% of Escherichia coli genes that lack experimental evidence of function. Nucleic Acids Res., 47, 2446–2454.

53. Santos-Zavaleta, A., Salgado, H., Gama-Castro, S., Sánchez-Pérez, M., Gómez-Romero, L., Ledezma-Tejeida, D., García-Sotelo, J.S., Alquicira-Hernández, K., Muñiz-Rascado, L.J., Peña-Loredo, P., et al. (2019) RegulonDB v 10.5: tackling challenges to unify classic and high throughput knowledge of gene regulation in E. coli K-12. Nucleic Acids Res., 47, D212–D220.

54. Keseler, I.M., Mackie, A., Santos-Zavaleta, A., Billington, R., Bonavides-Martínez, C., Caspi, R., Fulcher, C., Gama-Castro, S., Kothari, A., Krummenacker, M., et al. (2017) The EcoCyc database: reflecting new knowledge about Escherichia coli K-12. Nucleic Acids Res., 45, D543–D550.

55. Ishihama, A., Shimada, T. and Yamazaki, Y. (2016) Transcription profile ofEscherichia coli: genomic SELEX search for regulatory targets of transcription factors. Nucleic Acids Research, 44, 2058–2074.

56. Shimada, T., Fujita, N., Yamamoto, K. and Ishihama, A. (2009) Genomic SELEX for the genome-wide search of regulation targets by transcription factors: SELEX-clos and SELEX-chip procedures. 2009 International Symposium on Micro-NanoMechatronics and Human Science, 10.1109/mhs.2009.5351954.

57. Sévin, D.C., Fuhrer, T., Zamboni, N. and Sauer, U. (2017) Nontargeted in vitro metabolomics for high-throughput identification of novel enzymes in Escherichia coli. Nat. Methods, 14, 187–194.

58. Maeda, Y., Lin, C.-Y., Ishida, Y., Inouye, M., Yamaguchi, Y. and Phadtare, S. (2017) Characterization of YjjJ toxin of Escherichia coli. FEMS Microbiol. Lett., 364.

59. Herzberg, M., Kaye, I.K., Peti, W. and Wood, T.K. (2006) YdgG (TqsA) Controls Biofilm Formation in Escherichia coli K-12 through Autoinducer 2 Transport. Journal of Bacteriology, 188, 587–598.

60. Ibáñez de Aldecoa, A.L., Zafra, O. and González-Pastor, J.E. (2017) Mechanisms and Regulation of Extracellular DNA Release and Its Biological Roles in Microbial Communities. Front. Microbiol., 8, 1390.

