## Supplemental materials for "Unraveling the functions of uncharacterized transcription factors in *Escherichia coli* using ChIP-exo"

**Dataset S1** The strains used in this study

**Dataset S2** The conditions for genome-wide experiments

**Dataset S3** The statistics for ChIP-exo experiments

**Dataset S4** The genome-wide binding sites of candidate TFs in *E. coli* K-12 MG1655

**Dataset S5** The dataset of gene expression (TPM)

**Dataset S6** The differentially expressed genes of representative candidate TFs in *E. coli* K-12 MG1655

**Dataset S7** The homology analysis of candidate TFs using a library of Hidden Markov models

### Supplementary material

#### Deciphering regulatory targets of candidate transcription factors

***Local regulator (type II), YdhB***. We identified 29 genome-wide binding sites of YdhB in *E. coli* K-12 MG1655. There is one divergent binding peak among them located between gene *ydhB* and *ydhC* (**Supplementary Figure 5**). *ydhB* encodes the putative LysR-type DNA-binding transcriptional regulator, and *ydhC* encodes the putative transporter YdhC, which has been implicated in arabinose efflux (Koita & Rao, 2012). It is likely that YdhB represses transcription from the ydhC promoter and also interferes with its own transcription.

***Local regulator (type II), YneJ***. We identified 8 genome-wide binding sites of YneJ in *E. coli* K-12 MG1655 (**Supplementary Figure 6**). We found that YneJ directly controls the gene *sad* encoding succinate-semialdehyde dehydrogenase Sad. YneJ binds to the promoter of the operon *sad-yneH-yneG*, repressing gene expression when YneJ is present. However, the expression of the operon *sad-yneH-yneG* is activated during putrescine utilization (Rodionova et al., n.d.). Thus, our results showed that YneJ is involved in putrescine utilization.

***Local regulator (type II), YjhI***. We identified 5 genome-wide binding sites of YjhI in *E. coli* K-12 MG1655. We observed that YjhI has a binding site located at the upstream of operon *yjhIHG* (**Supplementary Figure 7**). The gene *yjhI* encodes the putative DNA-binding transcriptional regulator YjhI, *yjhH* encodes putative 2-dehydro-3-deoxy-Dpentonate aldolase, and *yjhG* encodes D-xylonate dehydratase. This operon regulates the energy conversion between pyruvate and glycolaldehyde. Thus, we inferred that YjhI is likely to be related to the conversion between pyruvate and glycolaldehyde.

***Local regulator (type II), YfiE***. We identified 4 genome-wide binding sites of YfiE in *E. coli* K-12 MG1655. We found that YfiE has one divergent binding peak between genes *yfiE* and *eamB* (**Supplementary Figure 8**). The gene *yfiE* encodes the putative LysR-type DNA-binding transcriptional regulator YfiE, and *eamB* encodes the cysteine/O-acetylserine exporter EamB. Thus, it is likely that YfiE is in the control of a cysteine and O-acetylserine exporter.

***Singly-target regulator (type III), YnfL.*** We identified a single target gene for YnfL in *E. coli* K-12 MG1655. This single binding site was located between the genes *ynfL* and *ynfM* (**Supplementary Figure 9**). *ynfL* encodes the putative DNA-binding regulator, and *ynfM* encodes the putative transporter YnfM. A previous study reported that YnfM is implicated in arabinose efflux (Koita & Rao, 2012). Therefore, it is possible that YnfL is a single-target regulator that controls the transporter YnfM that is involved in arabinose efflux.

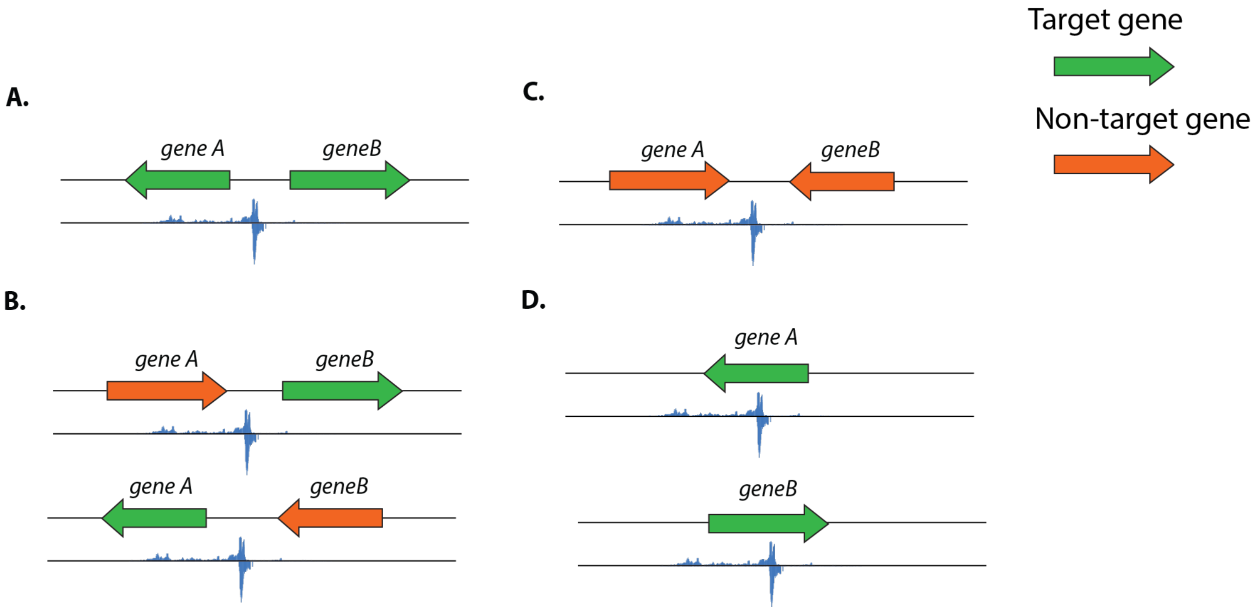


**Supplementary Figure 1 | The identification of target genes in peaking calling steps.**

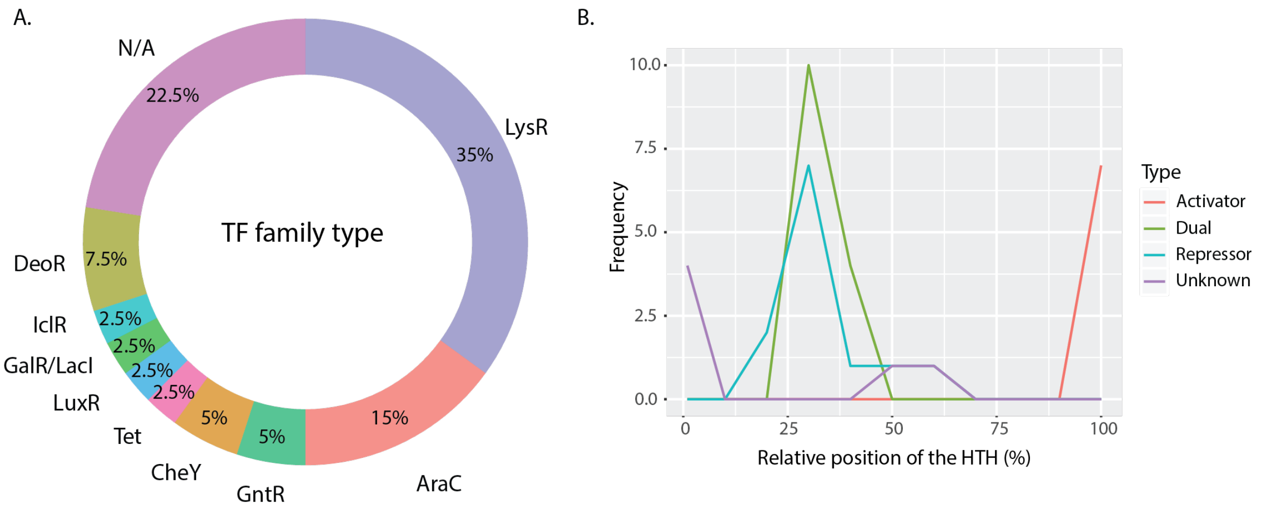


**Supplementary Figure 2 | The predicted family type for the 40 candidate TFs.**
**(A)** The classification of transcription factor family types for 40 candidate TFs. **(B)** Helix-Turn-Helix (HTH) distribution of 40 candidate TFs analyzed in this study. The distribution of the relative HTH location is shown for different subsets of the collection. On the x-axis, 0% represents the N-terminus and 100% represents the C-terminus of the protein. The y-axis is the number of candidate TFs. The four types of predicted results were colored in the legend. “Unknown” indicates that there is no prediction due to the lack of the information about HTH domain in the protein.

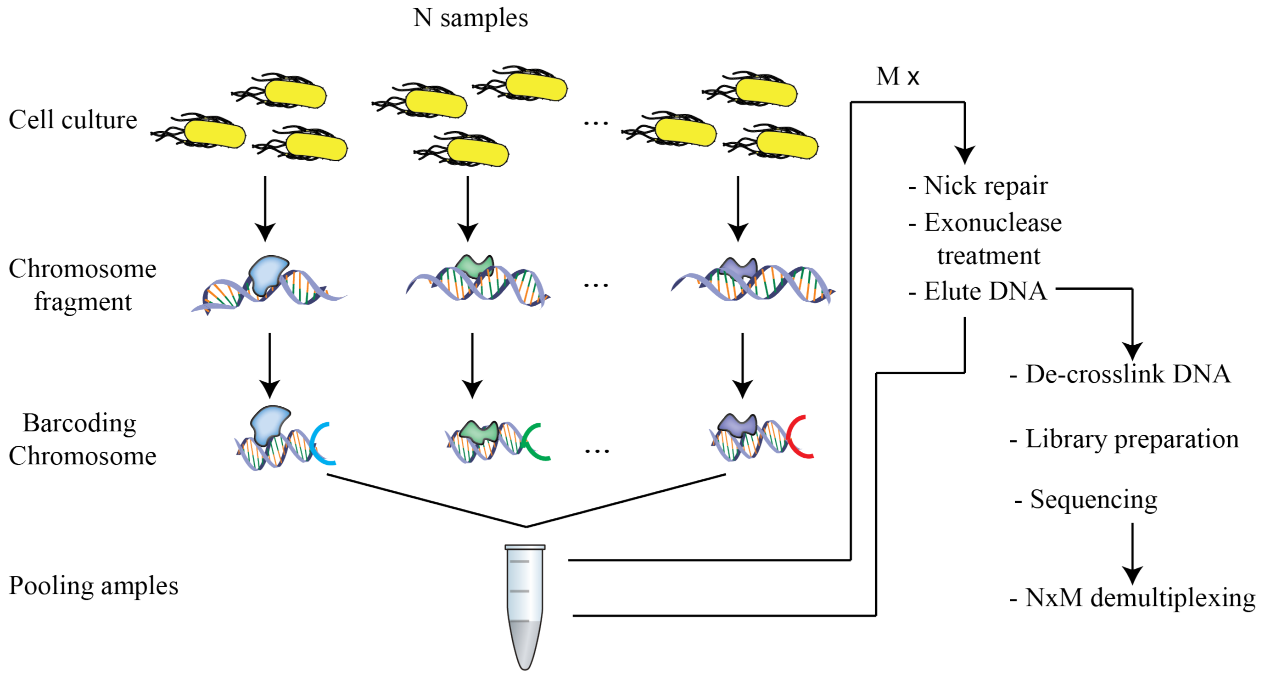


**Supplementary Figure 3 | The workflow of a multiplexed ChIP-exo system.** Cell cultures are crosslinked using formaldehyde. The complex (DNA bound to each candidate TF) is then extracted and fragmented using a sonicator. Sheared DNA fragments are immuno-precipitated with an antibody against myc-epitope. After ligation of the first adapter with specific indices, N fragmented samples have distinct indices for each other. Then they could be pooled and processed together (on-bead enzymatic reactions of the ChIP-exo method). The whole process could be repeated M times to generate N*M libraries. DNA recovered from the ChIP material can be directly amplified by PCR and deep-sequencing by Illumina next-generation sequencing technology.

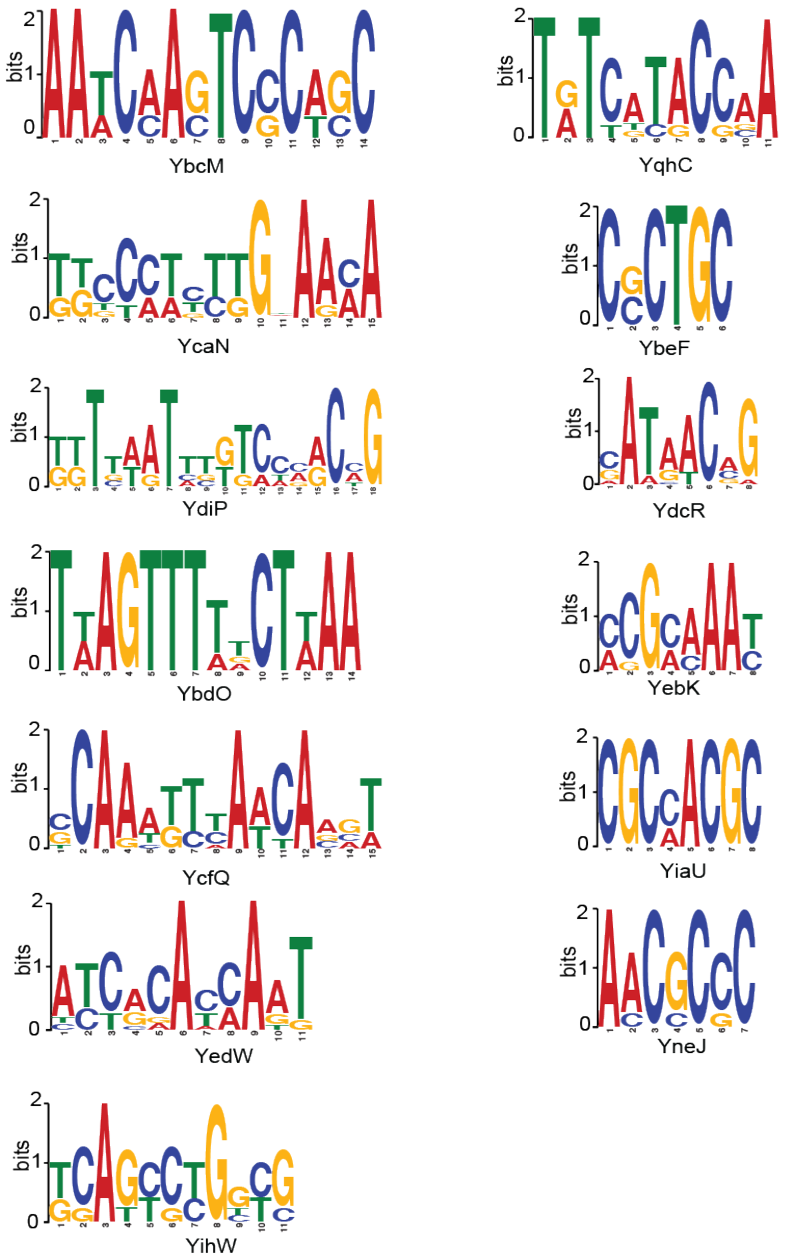


**Supplementary Figure 4 | The sequence motifs for candidate TFs.** The height of the letters (in bits on the y-axis) represents the degree of conservation at a given position within the aligned sequence set, with perfect conservation being 2 bits.

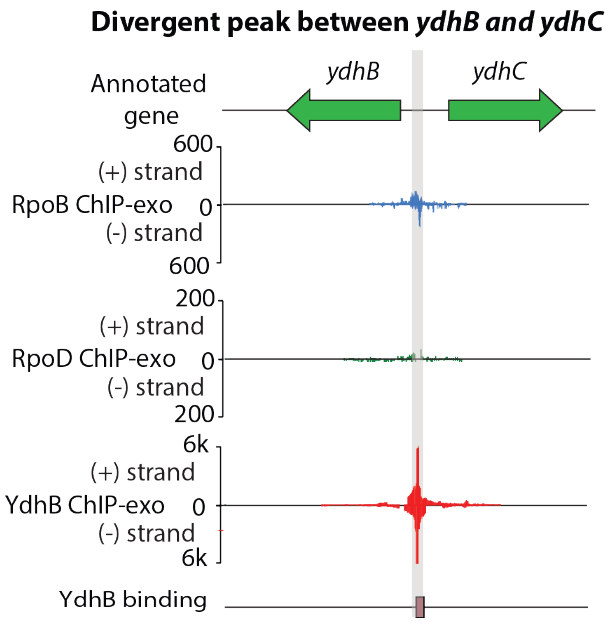


**Supplementary Figure 5 | Zoom-in divergent peak between *ydhB* and *ydhC*.**

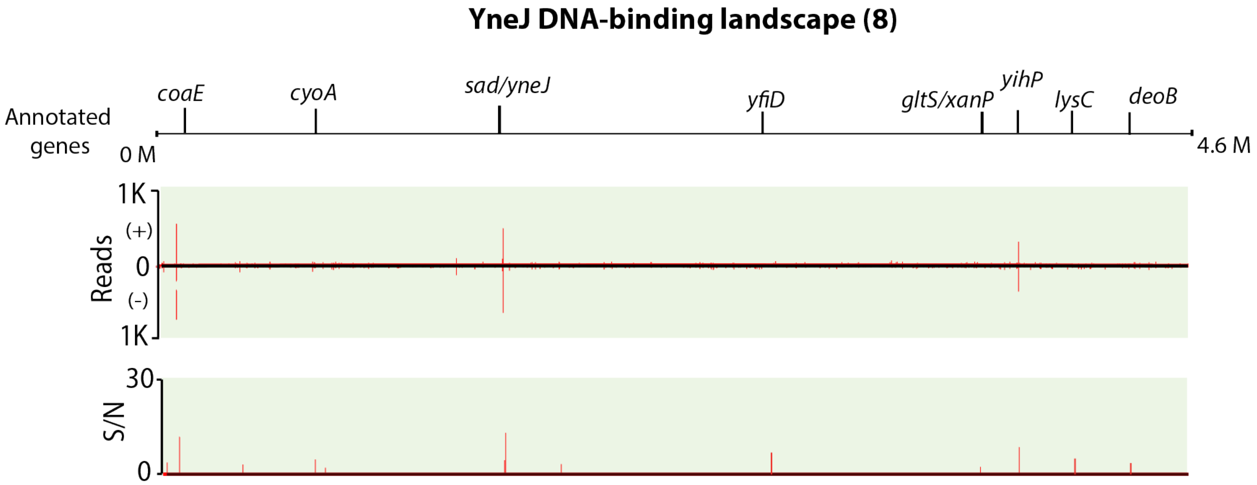


**Supplementary Figure 6 | Genome-wide binding of YneJ identified 8 binding events at the genome.**

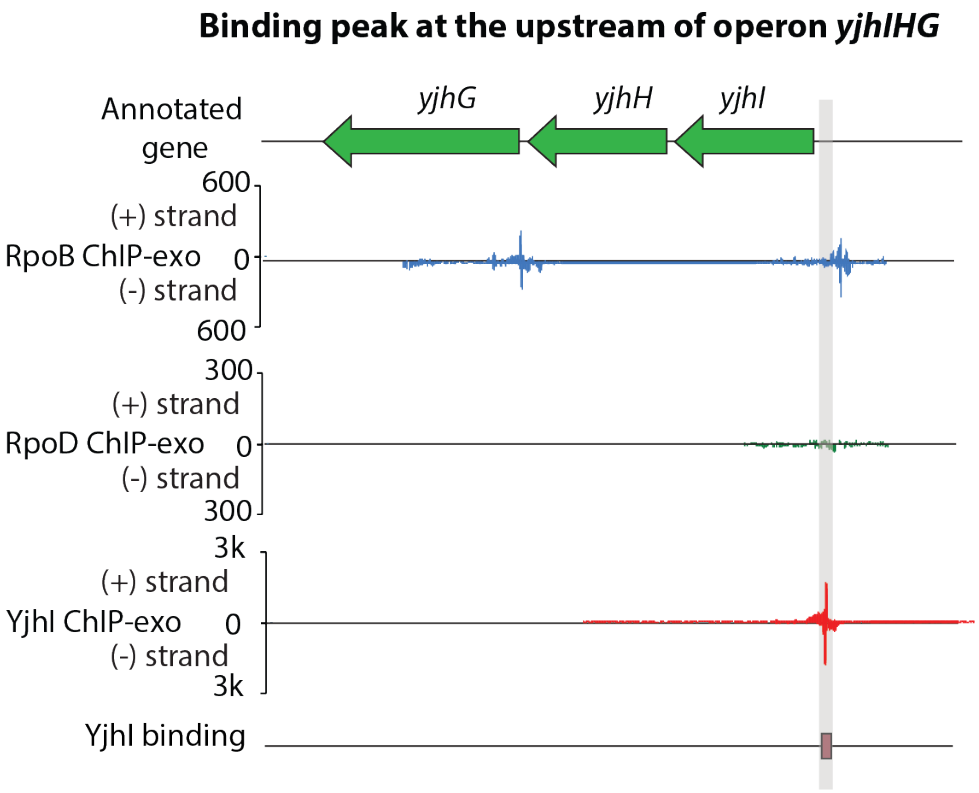


**Supplementary Figure 7 | Zoom-in binding peak at the upstream of operon *yjhIHG*.**

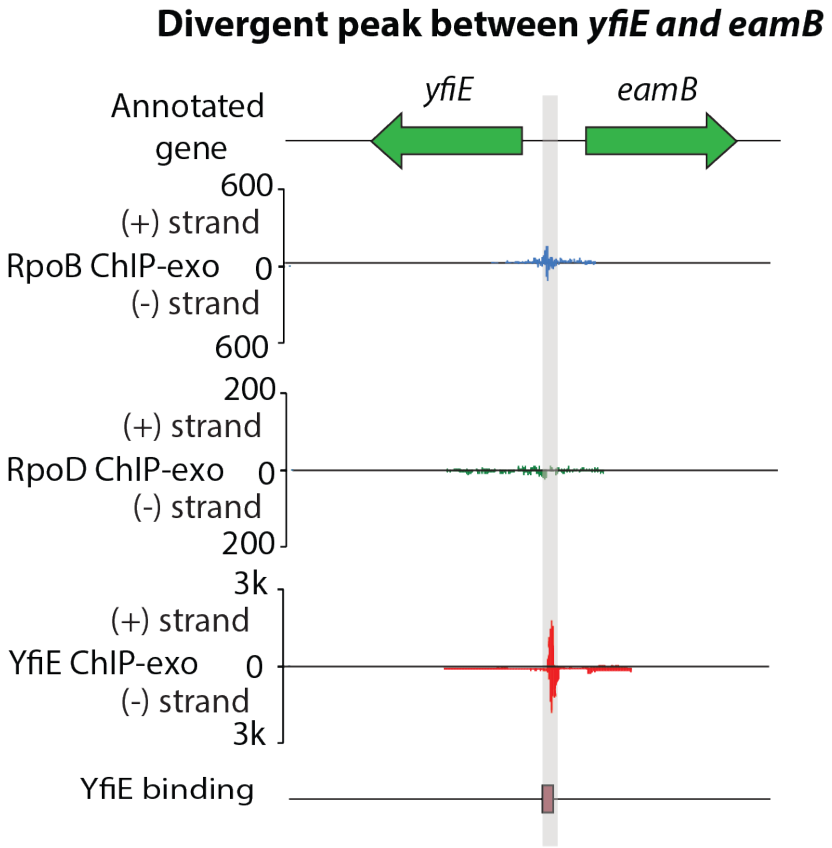


**Supplementary Figure 8 | Zoom-in divergent peak between *yfiE* and *eamB*.**

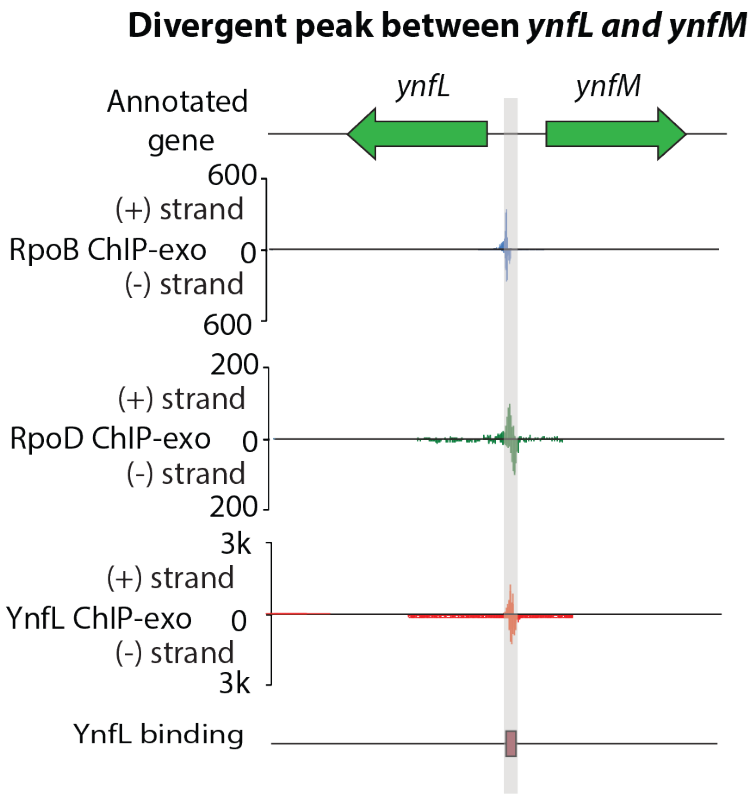


**Supplementary Figure 9 | Zoom-in divergent peak between *ynfL* and *ynfM*.**

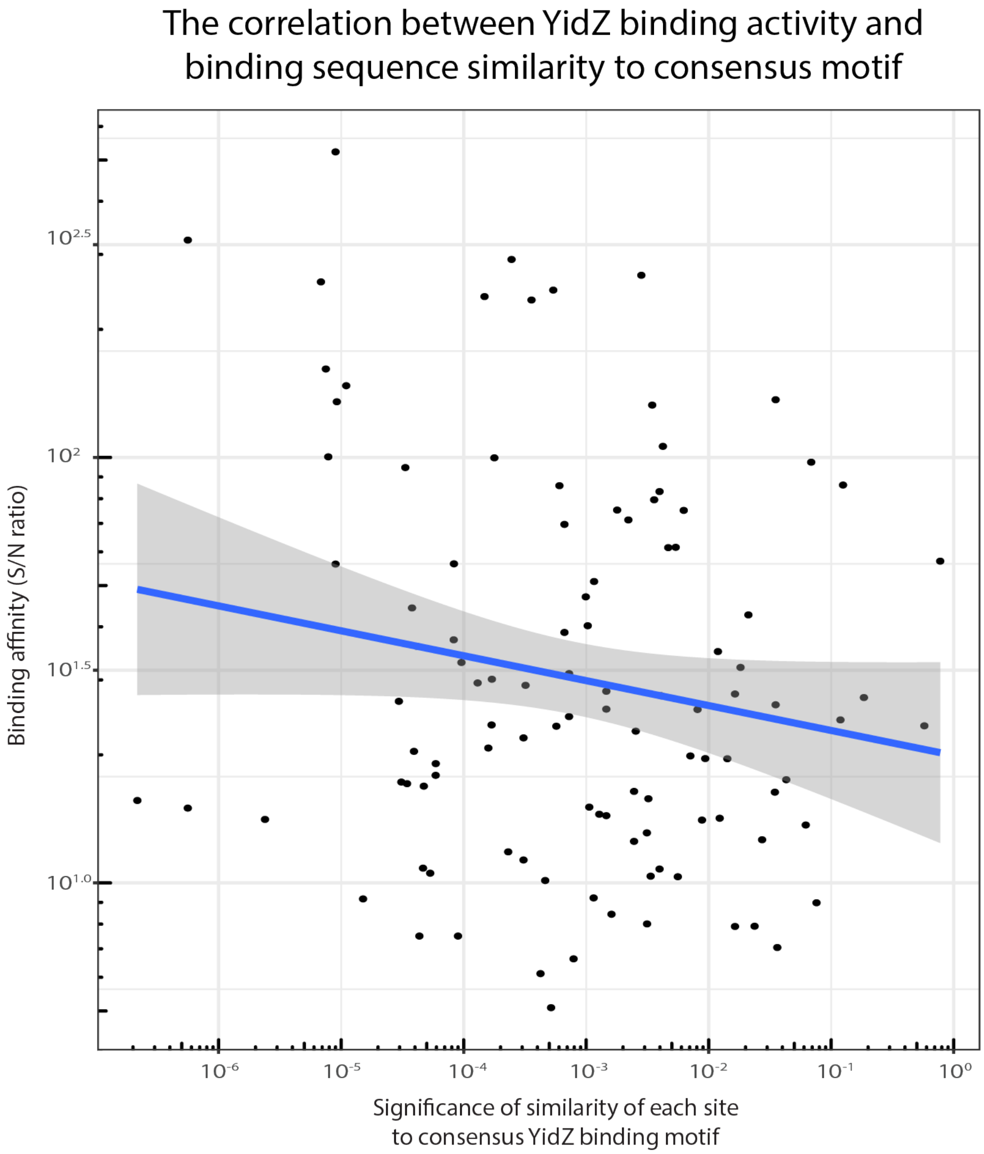


**Supplementary Figure 10 | Relationship between YidZ binding affinity and binding sequence similarity to consensus motif**. The signal-to-noise (S/N) ratio of each YidZ-binding peak was assigned to *in vivo* relative binding affinity of each site. The significance of similarity of each site to the consensus YidZ binding motif was represented by statistical significance (*p*-value) computed from the match score of the site with the position specific scoring matrix for the motif. More differences between the binding sequence and consensus motif resulted in increasingly lower binding affinity.

**Predicted YfeC structure**

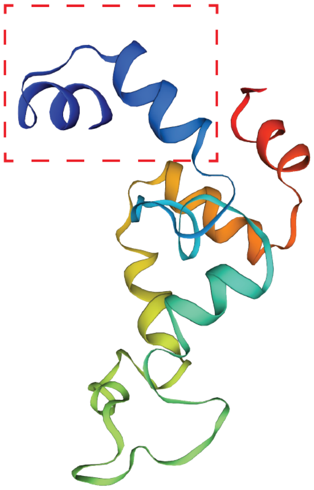

**Supplementary Figure 11 | Predicted 3D structure of YfeC (monomer with the range of AA 2-111).** The N-terminal region, labelled by a red dashed rectangle, is predicted as a DNA-binding domain by Hidden Markov Models.

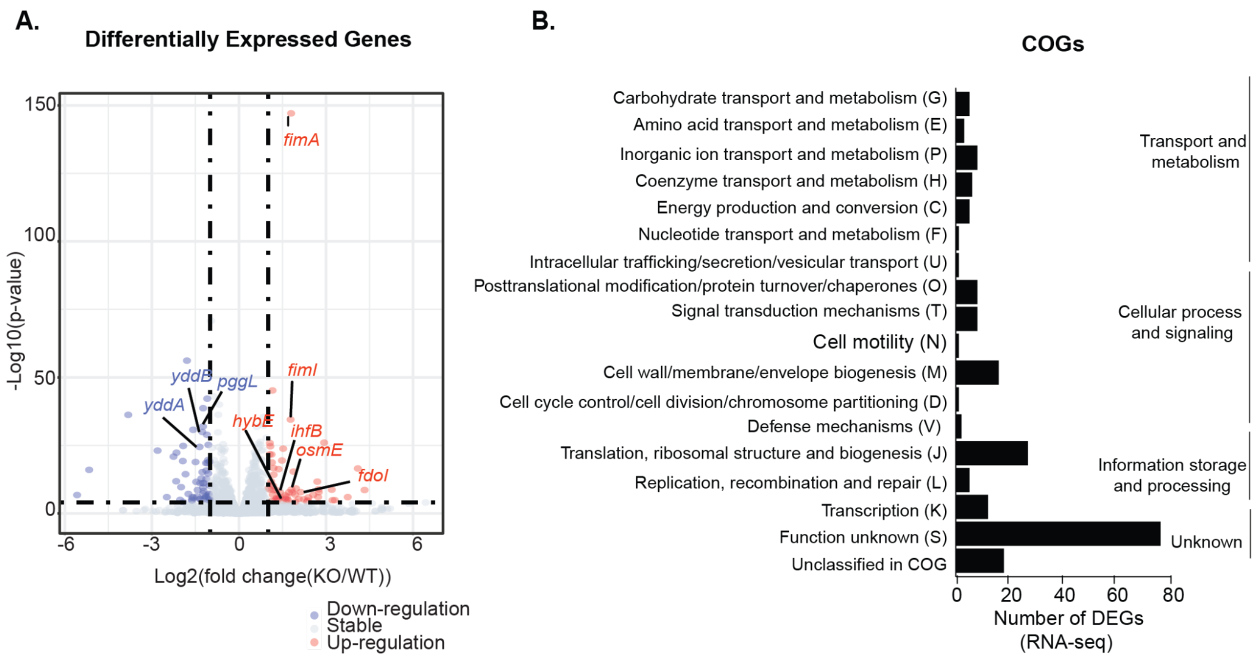


**Supplementary Figure 12 | Functional classification of differentially expressed genes in the *yfeC* deletion strain.** (A) 205 genes were differentially expressed after deletion of *yfeC* (cut-off value is $\log_2$ fold-change >=1, or <=-1, and adjust p-value < 0.05). (B) Functional classification of differentially expressed genes in the *yfeC* deletion strain. The top three clusters of functional groups were involved in transport and metabolism, cellular process and signaling, and information processing.

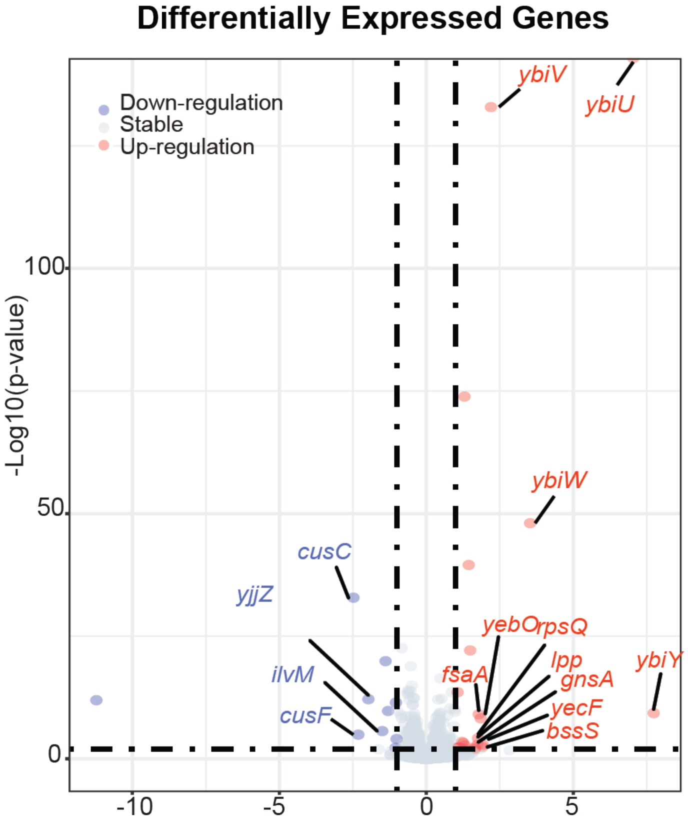


**Supplementary Figure 13 | The gene expression comparison between the wild type and the *yciT* deletion strains.** 46 genes were differentially expressed after deletion of *yciT* (cut-off value is $log_2$ fold-change >=1, or <=-1, and adjust p-value < 0.05).

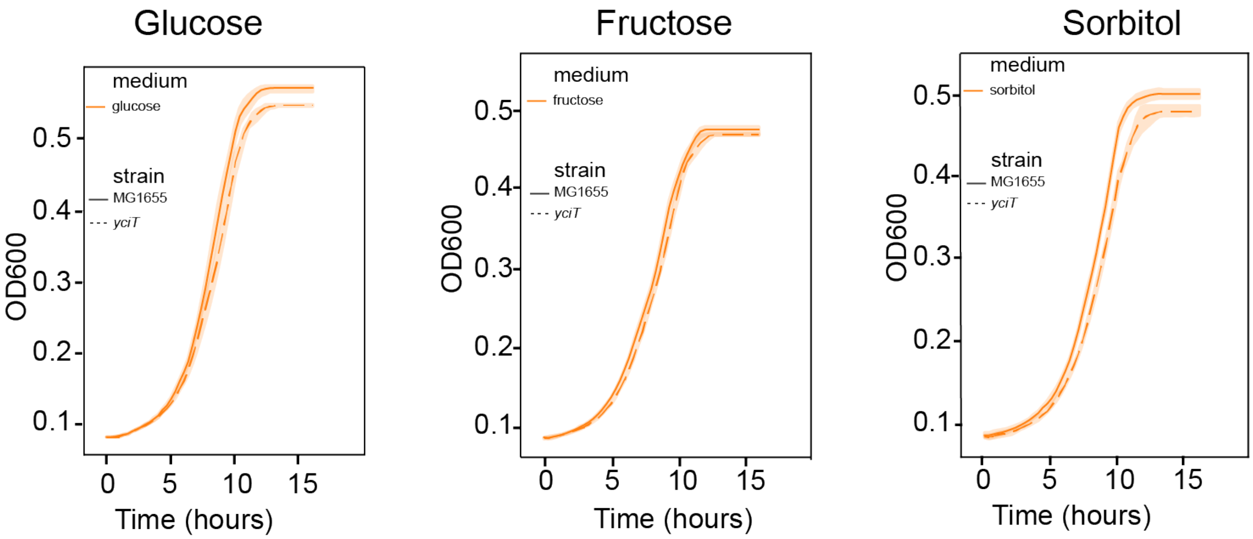


**Supplementary Figure 14 | The growth curve of wild type and *yciT* deletion strains at different carbon sources in the absence of osmotic stress (0.5 M NaCl).** Width of shaded bands represents standard deviation of the corresponding growth trajectory.

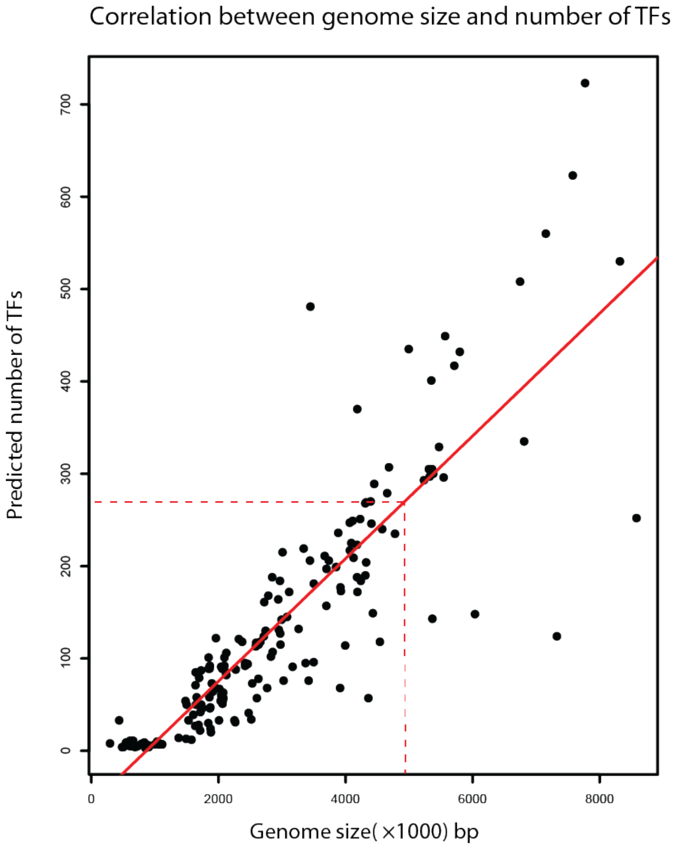


**Supplementary Figure 15 | The correlation between genome size and the number of TFs in the species.** The red line represents the linear correlation between them. Pearson correlation is 0.86. The red dashed line represents the relationship between *E. coli* K-12 MG1655 genome size (4.6Mbp) and estimated total number of TFs (280).

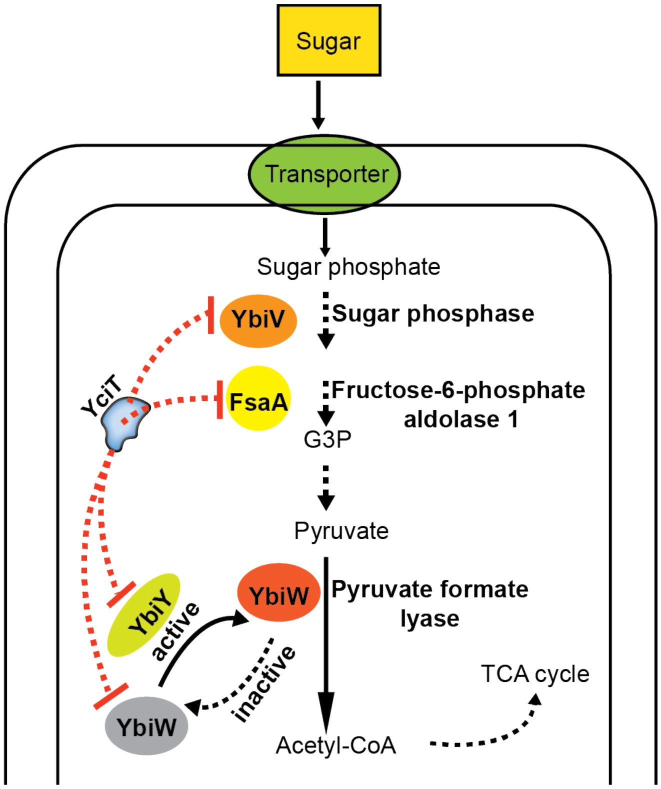


**Supplementary Figure 16 | The proposed pathway that is regulated by YciT**. When YciT is present (active) in *E. coli*, it represses the expression of target genes (*ybiV*, *ybiU*, *fsaA*, and *ybiY*).

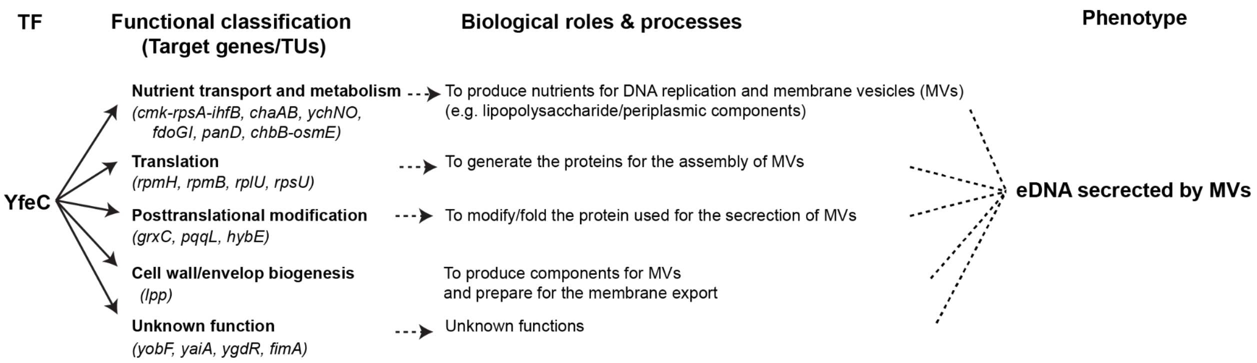


**Supplementary Figure 17 | YfeC regulatory networks regulate cellular functions.** Arrows with solid lines indicate that the regulation observed in this study. Arrows with dotted lines indicate that the regulation is not observed in this study.

**Supplementary Table 1** Comparison of structure and regulatory activities between candidate TFs and known TFs in *E. coli* K-12 MG1655.

| | Known TFs | | | Candidate TFs | | |
|---|---|---|---|---|---|---|
| Family Type | Regulatory Role | Average size of AAs | Physiological function | Predicted regulatory Role | Average size of AAs | Representatives |
| LysR | Dual | Varied | Amino acid biosynthesis | Dual | ~ 306 | YahB, YbdO |
| AraC | Activator | Varied | Virulence, sugar metabolism | Repressor | ~ 280 | YbcM |
| GntR | Repressor | ~ 246 | Carbon metabolism | Repressor | ~ 352 | YdcR |
| TetR | Repressor | ~ 210 | Tetracycline resistance | Repressor | ~ 210 | YcfQ |
| LuxR | Activator | ~ 220 | Biosynthesis and glycerol metabolism | Activator | ~ 200 | YhjB |
| GalR/LacI | Repressor | ~ 334 | Carbon source uptake | Repressor | ~ 332 | YcjW |
| IclR | Repressor | ~ 273 | Carbon source uptake | Repressor | ~ 262 | YjhI |
| DeoR | Repressor | ~ 257 | Sugar metabolism | Repressor | ~ 261 | YihW, YciT, YgbI |

N/A* indicates that there are no representative candidates for this condition.

**Supplementary Table 2** Protein sequence metadata and homology modeling statistics from SWISS-MODEL

| | | Source: UniProt | | | Source: SWISS-MODEL | | | |
|---|---|---|---|---|---|---|---|---|
| **Gene ID** | **Protein name** | **UniProt** | **DNA binding domain motif** | **DNA binding domain residues** | **PDB template** | **GMQE (Global Model Quality Estimate)** | **Oligomeric state prediction** | **QMEAN Z-score** |
| b3711 | YidZ | P31463 | H-T-H motif | 25 to 44 | 2esn.1.A | 0.50 | Dimer or Tetramer | -3.14 |
| b2398 | YfeC | P0AD37 | N/A | N/A | 2qww.1.A | 0.34 | Dimer | -3.17 |
| b1284 | YciT | P76034 | H-T-H motif | 18 to 37 | 2w48.1.A | 0.34 | Tetramer | -5.31 |
| b2735 | YgbI | P52598 | H-T-H motif | 20 to 39 | 2w48.1.A | 0.36 | Tetramer | -5.57 |
| b1320 | YcjW | P77615 | H-T-H motif | 5 to 24 | 1vpw.1.B | 0.67 | Dimer | -2.36 |
| b1434 | YdcN | P77626 | H-T-H motif | 23 to 42 | 1y9q.1.A | 0.75 | Dimer | -1.94 |

Note, N/A indicates that there is no available information in the UniProt.

**Supplementary Table 3** The target genes directly regulated by YfeC

| b_number | Transcriptional unit | Description/Function | COGs | log2(FoldChange) | padj |
|---|---|---|---|---|---|
| b3703 | *rpmH* | 50S ribosomal subunit protein L34 | Translation, ribosomal structure and biogenesis | 4.33079132 | 4.36E-08 |
| b3637 | *rpmB* | 50S ribosomal subunit protein L28 | Translation, ribosomal structure and biogenesis | 1.15981833 | 0.01940833 |
| b3186 | *rplU* | 50S ribosomal subunit protein L21 | Translation, ribosomal structure and biogenesis | 1.59875753 | 0.00000297 |
| b3065 | *rpsU* | 30S ribosomal subunit protein S21 | Translation, ribosomal structure and biogenesis | 1.93796714 | 0.00230352 |
| b2399 | *yfeCD* | putative transcription factor | Function unknown | 4.32546198 | 4.98E-133 |
| b1161 | *ycgFX* | blue light- and temperature-regulated antirepressor BluF | Function unknown | -1.6727558 | 0.01324743 |
| b3610 | *grxC* | reduced glutaredoxin 3 | Post-translational modification, protein turnover, chaperones | 1.65174882 | 0.00000821 |
| b1494 | *pqqL* | periplasmic metalloprotease | Post-translational modification, protein turnover, chaperones | -1.0556295 | 8.21E-24 |
| b2992 | *hybE* | hydrogenase 2-specific chaperone | Post-translational modification, protein turnover, chaperones | 1.51293908 | 0.00021498 |

**Supplementary Table 3** The target genes directly regulated by YfeC (con't)

| b_number | Transcriptional unit | Description/Function | COGs | log2(FoldChange) | padj |
|---|---|---|---|---|---|
| b0910 | *cmk-rps-ihfB* | cytidylate kinase | Nucleotide transport and metabolism | 1.20143761 | 3.07E-09 |
| b1216 | *chaAB* | Na+/K+:H+ antiporter ChaA | Inorganic ion transport and metabolism | 1.55342508 | 0.01017597 |
| b1219 | *ychNO* | DsrE/F sulfur relay family protein YchN | Inorganic ion transport and metabolism | -1.3343157 | 7.36E-12 |
| b3894 | *fdoGI* | formate dehydrogenase O subunit α | Energy production and conversion | 1.16191449 | 0.00386399 |
| b0131 | *panD* | aspartate 1-decarboxylase proenzyme | Coenzyme transport and metabolism | -1.3441817 | 0.0000505 |
| b4317 | *fimAICD* | outer membrane protein; export and assembly of type 1 fimbriae | Cell wall/membrane/envelope biogenesis | 1.05862358 | 1.91E-22 |
| b1677 | *lpp* | murein lipoprotein | Cell wall/membrane/envelope biogenesis | 1.7414382 | 0.00345401 |
| b1738 | *chbB-osmE* | N,N'-diacetylchitobiose-specific PTS enzyme IIB component | Carbohydrate transport and metabolism | 1.80868592 | 1.81E-09 |
| b1824 | *yobF* | small protein involved in stress response | Function unknown | 2.40258368 | 0.00952181 |
| b0389 | *yaiA* | protein YaiA | Function unknown | 1.36869395 | 0.015577 |
| b2833 | *ygdR* | predicted protein | Function unknown | 1.53268718 | 0.00741777 |

### References

Koita, K., & Rao, C. V. (2012). Identification and analysis of the putative pentose sugar efflux transporters in Escherichia coli. *PloS One*, *7*(8), e43700.

Lau, M. E., Loughman, J. A., & Hunstad, D. A. (2012). YbcL of uropathogenic Escherichia coli suppresses transepithelial neutrophil migration. *Infection and Immunity*, *80*(12), 4123–4132.

Rodionova, I. A., Gao, Y., Sastry, A., Monk, J., Wong, N., Szubin, R., Lim, H., Zhang, Z., Saier, M. H., & Palsson, B. (n.d.). *PtrR (YneJ) is a novel E. coli transcription factor regulating the putrescine stress response and glutamate utilization*. https://doi.org/10.1101/2020.04.27.065417

Serre, L., Pereira de Jesus, K., Zelwer, C., Bureaud, N., Schoentgen, F., & Bénédetti, H. (2001). Crystal structures of YBHB and YBCL from Escherichia coli, two bacterial homologues to a Raf kinase inhibitor protein. *Journal of Molecular Biology*, *310*(3), 617–634.
